# A breath of fresh air: comparative evaluation of passive versus active airborne eDNA sampling strategies

**DOI:** 10.1101/2025.03.26.645491

**Authors:** Hugo Jager, Krijn B. Trimbos, Jan-Maarten Luursema, Adrianus G.C.L. Speksnijder, Kathryn A. Stewart

## Abstract

Terrestrial biodiversity’s rapid decline demands expansion of high-resolution biomonitoring to support science-based policy. Environmental DNA (eDNA) analysis has proven effective in aquatic systems but remains underexplored for terrestrial habitats, partly due to point-source sampling bias and difficulty in upscaling. Air has emerged as a promising substrate yet most studies use active samplers which tend to be expensive, bulky, and require power. We used a newly developed, inexpensive, reusable, and easy-to-use passive airborne eDNA sampler (Nutshell eDNA sampler) to capture eDNA suspended in air across time (6 to 96 hours) within Rotterdam Zoo, the Netherlands. Its performance was compared to two commonly-used active airborne eDNA samplers for vertebrate diversity detection. In total, 88 species were detected, including 24 zoo residents. The Nutshell eDNA sampler was the most effective at detecting zoo residents, surpassing active samplers in species richness within 48 hours and continuing to accumulate new species beyond 96 hours, including detections of both patchy and singleton signals. It also detected the furthest species signal (515 m). Zoo airborne eDNA further demonstrated a positive correlation to species total biomass, suggesting larger vertebrates release proportionately more DNA into the air. Our findings indicate that for long, unsupervised biomonitoring, passive airborne eDNA sampling presents a promising approach for assessing vertebrate communities and putatively reduces detection noise in stochastic air eDNA signals. While deeper investigations into airborne eDNA sampling strategies are needed, passive methods can offer a much needed logistically flexible, low-maintenance approach compared to short-burst collection strategies employed by many ac-tive samplers.

## Introduction

Global biodiversity is essential for ecosystem resilience and critical services such as food security and medicine [1,2]. However, anthropogenic pressures such as climate change, habitat destruction, and globalisation threaten biodiversity. This necessitates high-resolution biomonitoring that can be quickly upscaled to guide conservation efforts [3,4].

While most biodiversity monitoring relies on conventional approaches (e.g. morpho-logical surveys), they are often time-consuming, require expert taxonomic knowledge, can be invasive to the species, and fail to document rare species [5]. Environmental DNA (eDNA) approaches, which collect genetic material directly from the environment rather than indi-viduals [5–7], have recently gained widespread use as an alternative or complimentary method [4]. These approaches are non-invasive, highly sensitive, cost effective, and capable of detecting taxa across the tree of life while reducing observer bias and enabling large-scale monitoring [8,9]. In many cases, eDNA collection outperforms conventional methods [10,11]. While eDNA sampling from water is well-established [12–14], terrestrial applications remain underdeveloped partly due to challenges in identifying optimal substrates [15], particularly for biomonitoring upscaling [16]. Various terrestrial substrates, including soil, plants and spider webs (among others) have been explored but often yield limited taxonomic coverage and rely on point-source samples that may fail to give a comprehensive and representative estimate of biodiversity [17–20].

Airborne eDNA sampling offers a novel approach for terrestrial biodiversity monitor-ing. Air contains *bioaerosols*, or microscopic particles of organic origin suspended in air.

Bioaerosols tend to be rich in genetic material that can originate from all domains of life [21]. Researchers [22] first demonstrated animal DNA detection in air using active (powered) air pumps, with subsequent studies confirming this method’s efficacy across various environ-ments, including zoos and natural habitats [23–25]. Recent research has detected insects, birds, frogs, and mammals in field conditions, highlighting the potential of airborne eDNA for large-scale biodiversity monitoring across the tree of life [7,21,26].

Most eDNA studies focusing on air collection to date rely on active samplers, which, despite their effectiveness, are often costly, bulky, and require power sources, limiting their use in remote areas [27]. For the most part, they also rely heavily on short-burst point-sampling due to logistical (e.g., battery life and personnel) constraints. Additionally, budget-ary and personnel constraints can limit the scale at which active sampling is applied. Passive air eDNA sampling, which relies on natural air movement to deposit DNA-containing particles (like skin cells, hair, spores, pollen, or fecal dust) onto a collection surface, on the other hand, potentially provides a cheaper and more mobile alternative, with promising results for detecting diverse taxa [28–30]. Recent studies using passive approaches have repurposed existing (e.g. dust collectors [28], spider webs [31]) or rudimentary tools (e.g. buckets of wa-ter [30]) with great success. However, expanding air eDNA collections may require more lo-gistically and technologically flexible solutions. Additionally, questions remain regarding the sensitivity and spatial resolution of passive approaches compared to active methods, espe-cially given evidence of long-range airborne eDNA dispersal [23] and continuous accumula-tion over time [30]. Thus, not only are passive-active eDNA approach comparisons neces-sary, understanding airborne eDNA’s spatial and temporal dynamics is also crucial for opti-mizing and choosing the appropriate terrestrial sampling strategies [31].

Comparing passive to active eDNA sampling strategies have only just recently been conducted for aquatic eDNA collection strategies [32, 33], but none to date have been at-tempted with eDNA collections of air. Our study thus compared two general strategies for air eDNA collection: passive and active approaches in which we evaluated the efficacy and sen-sitivity of a newly-designed passive airborne eDNA sampler (the Nutshell eDNA Sampler; Supplementary Information-Methods; Fig. S1 & Fig S2 for sampler details). Our sampling strategy comparison employs two, commonly used, active air eDNA samplers (Pollensniffer and Coriolis Micro) as a reference in Rotterdam Zoo, The Netherlands. The Nutshell eDNA sampler, designed to enhance DNA capture efficiency by directing airflow over a protected DNA-binding filter, aims to mitigate the limitations of previous passive methods that rely on surface settlement rather than via air currents themselves [28, 30, 34–35]. Addressing key needs for upscaling biomonitoring via air eDNA, the Nutshell eDNA sampler is an open source design which is mostly 3D printed, can be easily assembled, and is designed for re-use, affordability, and sustainability (see Supplementary Information). To test how a sampling strategy employing the Nutshell eDNA sampler performed, we compared it to two commonly used active collection approaches, assessing species detection rates, expected zoo verte-brate species recovery, airborne eDNA dispersion distances, accuracy in detecting known zoo residents, and a potential correlation between species biomass and airborne eDNA abundance. Additionally, we looked at species richness over time to determine the optimal deployment duration for our passive approach compared to the typical short-burst approach of active sampling.

## MATERIALS AND METHODS

### Study site

We collected air samples at Rotterdam Zoo (Diergaarde Blijdorp, the Netherlands; Figure 1), a 32-acre zoo wherein sampling took place at five outdoor points and one indoor point. Sur-rounded by residential areas and water bodies, the zoo houses diverse avian and mammal-ian species in spacious outdoor enclosures. Besides listed zoo animals, expected eDNA sources include food, visitors, their pets, and nearby wildlife.

**Figure 1:**
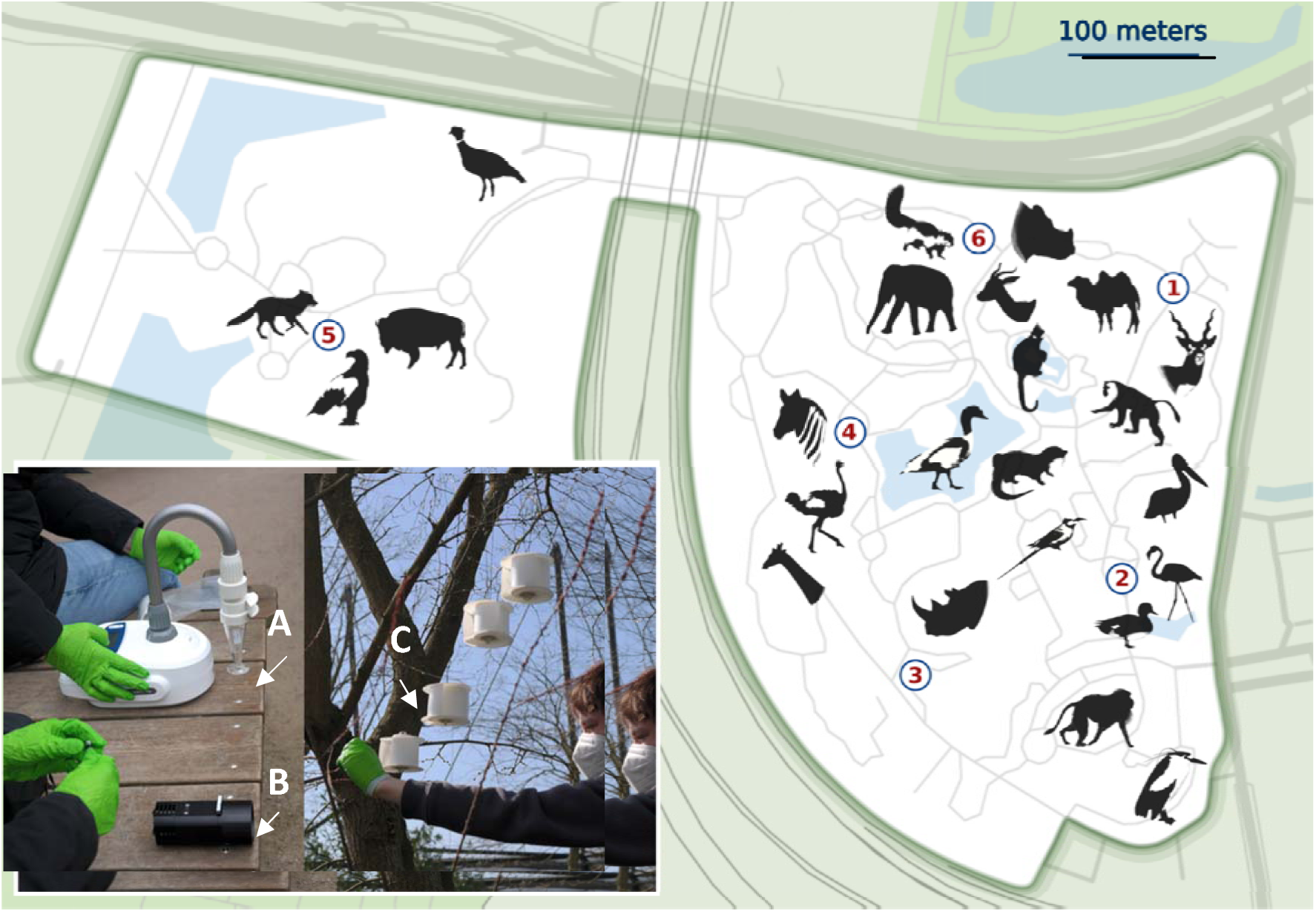
Map of Rotterdam Zoo (the Netherlands) including the 6 sampling locations (numbered circles). Location 6 represents an indoor site. Photo insert depicts sampling setup of the Coriolis Micro (**A**), Pollensniffer (**B**), and the passive Nutshell eDNA samplers (subject is H. Jager) (**C**).

### Airborne eDNA sampling

Between March 25 and April 12 2024, we collected air samples at six locations over three 5-day sampling campaigns (Supplementary Information, Table S1). Locations were selected based on: 1) secure placement of passive airborne eDNA samplers out of visitor’s reach, 2) sufficient distance between sampling points to avoid redundant detections, and 3) proximity to at least four enclosures.

To test the performance of the newly developed passive Nutshell eDNA sampler with active samplers as a reference, three types of air sampling device were used, each varying in mobility, cost, and flow-rate: two active samplers - the Pollensniffer (∼8 L/min) which collects eDNA on Vaseline coated slides [36], the Coriolis Micro (∼100 L/min) which collects eDNA in collection liquid [37], and the passive Nutshell eDNA sampler [38] which relies on external air movement and collects eDNA on glass-fibre filters. To minimize disruption to animals in exhibit in consultation with zoo staff, while acting in accordance with suggested best sampling practices by the manufacturers, both active samplers were operated simultaneously for 10 minutes per sampling event. The passive air DNA sampling strategy used here (Nutshell eDNA sampler) allows for longer deployment, which increases the potential amount of air DNA captured. However, we had no *a-priori* expectation for how well the sampler will collect air eDNA and thus deployed the Nutshell eDNA sampler across 5 temporal durations. To test for optimal deployment time (the time at which the maximum number of species would be captured) for our passive approach, five Nutshell eDNA samplers were deployed per sampling location, which were subsequently collected at 6, 24, 48, 72, and 96 hours. This allowed us to create species saturation curves. We deployed the passive samplers at ∼2.5 m height in trees at each location (including the indoor location, also see Figure 1) on the first day of each sampling campaign, and took 1 m-high air samples with both active samplers at each site (replicating common methodological approaches [33][34]). Each sampling campaign (the Nutshell eDNA, Pollensniffer and Coriolis Micro) was replicated three times (consecutive weeks) to attain triplicate measures for each active and passive sampler per site. All equipment was sterilized between uses in a 30% bleach solution, followed by ultrapure water rinsing. Passive Nutshell eDNA samples were collected on 47 mm diameter 0.7 µm glass fibre filters, stored in 2 ml Eppendorf tubes. The Coriolis Micro samples were collected in 4 mL PBS-filled cones, sealed immediately after sampling. The Pollensniffer samples were collected on Vaseline-coated plastic slides and stored in a sealed cassette case. Samples were handled with facemasks and nitrile gloves and transported on ice before storage the same day at -20 °C until DNA extraction.

For each campaign, field negative controls were also collected to account for any potential DNA contamination in the collection media or on the samplers themselves prior to sampling at the zoo: glass fibre filters placed inside the sampler but without being hung (Nutshell eDNA Sampler), PBS solution placed within the sampling cup (Coriolis Micro), and a Vaseline-coated slide (Pollensniffer). In total for six locations and three sampling replication events, 90 Nutshell eDNA samples (18 x 5 time intervals at 6, 24, 48, 72 and 96 hours), 18 Pollensniffer samples, 18 Coriolis Micro samples, and three field negatives per device were obtained across the three weeks.

### DNA extraction

For extraction, we fit each sample in a 2 mL Eppendorf tube: Nutshell eDNA sampler filters were cut into small pieces in a sterilized, eDNA dedicated fume-hood, the Pollensniffer slides were removed from the cassettes and folded with the exposed side facing inwards, the 4mL Coriolis Micro PBL samples were transferred to an Eppendorf tube, centrifuged for two minutes at 10,000 rpm, thereafter supernatant removed and pellets retained. We extracted DNA for all samples following the protocol of [39]. Samples were incubated overnight in a 56°C lysis buffer for 14.5 hours. QIAshredder (QIAGEN) was then used to release lysed cellular content from the sample substrates, which after centrifuging for three minutes at 11,000 rpm resulted in two 390 µL tubes per sample. We vortexed samples and added 700 µL AL:EtOH (1:1) to each tube. Finally, we eluted DNA in 100 µL AE buffer. All samples were stored at - 20 °C until metabarcoding.

### DNA metabarcoding

We used two vertebrate primer sets for metabarcoding: 16S rRNA primer (16Smam1/16Smam2) targeting mammals (∼95 bp) [40], and 12S rRNA (12S05 forward /12S05 reverse) targeting vertebrates (∼97 bp) [41]. All primers had 5’-overhang adapters for Nextera XT (Illumina) indexing. A 1:100 dilution of harbor porpoise (*Phocoena phocoena*) DNA (∼50 ng/µL; Naturalis Biodiversity Center, Netherlands) served as a positive PCR con-trol, with multiple negative controls to detect contamination.

For the 16S primer set, 20 µL reactions consisted of 4 µL DNA template, 10 µL TaqMan™ Environmental Master Mix 2.0, 1 µL of each primer (10 µM), 3 µL RNase-free wa-ter, and 1 µL human blocker (60 µM) [42]. The thermal cycling profile was 95 °C for 10 min-utes, followed by 40 cycles of 95 °C for 12 s, 59 °C for 30 s, 70 °C for 25 s, and a final exten-sion time of 72 °C for 7 minutes. The 12S reactions followed the same protocol but omitted the human blocker [43] and had a cycling profile of 94 °C for 30 s, 59 °C for 45 s, and 72 °C for 60 s. Three PCR replicates were performed for all 131 DNA extracts, controls, and both primer sets.

PCR products were visualized on 2% agarose gels (Invitrogen) with SYBR™ Safe alongside a low-range quantitative DNA ladder. Three technical replicates (10 µL each) were pooled per DNA extract. Amplicon pools were purified using NucleoMag NGS Clean-up and Size Select beads (Mackerey-Nagel) at a 1.6× bead-to-amplicon ratio. Nextera XT (Illumina) indices were ligated in a second PCR (20 µL), consisting of 2 µL amplicon pool, 10 µL TaqMan™ Environmental Master Mix 2.0, 1.5 µL of both N- and S-indices, and 5 µL RNase-free water. The thermal profile was 95 °C for 10 min, 10 cycles of 95 °C for 15 s, 55 °C for 30 s, 72 °C for 1 min, and a final extension at 72 °C for 5 min.

Amplicon pool concentrations were measured using the 4200 Tapestation System (Agilent Technologies) and pooled equimolarly using the OT-2 Robot (Opentrons). The final pool was purified using a 1.2× bead-to-amplicon ratio and quantified on the 4200 Tapesta-tion. Libraries were sequenced at Macrogen Europe (Netherlands) on an Illumina MiSeq (v3 chemistry, 150 bp paired-end), targeting 50,000 reads per sample.

### Data preparation

Quality assurance was performed separately for each primer set. Illumina adaptors and primers were trimmed using Cutadapt v4.9 [44], and reads were processed into amplicon se-quence variants (ASVs) using DADA2 [45] in R [46]. Filtering parameters included maxEE = 2 and minLen = 50, removing reads with >2 expected errors or <50 bp. Dereplication and error estimation followed, using DADA2 to calculate ASV counts. We merged reads, re-moved chimeras, retaining only ASVs between 80 -106 bp for 12S, and 83 - 118 bp for 16S.

Taxonomic identification was performed using BLASTn against GenBank 12S and 16S databases via a custom pipeline on Naturalis Biodiversity Center’s OpenStack GALAXY platform [47]. A weighted LCA algorithm applied cutoffs of 85% query coverage, 98% iden-tity, and a minimum score of 170 [48]. ASVs identified as human, unidentified, or detected in controls were removed. To minimize contamination, read counts in the negative PCR con-trols were subtracted from Nutshell eDNA sampler reads for their respective weeks and from all reads for Coriolis Micro and Pollensniffer. ASVs of the same species were grouped for both primer sets.

Read counts from both primers were combined into an ASV table summarizing total species read counts per sample. Separate datasets were created for each eDNA sampling strategy (active: Pollensniffer, Coriolis Micro; passive: Nutshell eDNA Sampler), summing species-specific reads across all samples for each method. Identified species were cross-referenced with a detailed Rotterdam Zoo inventory (N = 266), including those housed off-exhibit.

### Quantification and statistical analyses

We assessed normality of our data by using Shapiro-Wilk and Kolmogorov-Smirnof (KS) tests in R 4.4.0. We found all data met normality expectation with the exception of our sepa-rate dataset for zoo species sequencing reads, specifically for the Nutshell eDNA sampler between the outdoor sampling locations (p = 0.046), and total biomass (p > 0.05), both of which had to be log-transformed. To compare species richness detected by the three air-borne eDNA sampling strategies, we performed ANOVA and Tukey’s HSD tests. We further used a two-way ANOVA to check for significant effects (and their interaction) with replicate sampling weeks on sampling strategies.

Species richness accumulation over time was visualized using *vegan* v.2.6.6.1 and *ggplot2* [49,50]. All Nutshell eDNA samples (n = 75) were analyzed across outdoor sites (five locations in triplicate, each with five samplers collected at 6, 24, 48, 72, and 96 hours). Two species diversity saturation curves were generated: one for all detected species and one for listed zoo species only, visualizing for each time point the mean and SD of the total detected species richness per outdoor sampling location across the three sampling campaigns. For active sampler strategies (Pollensniffer and Coriolis Micro), total detected species were plot-ted per replication. For the active strategies, we report static values since the active samplers were not tested over a time series (6, 24, 48, 72, and 96 hours like the passive samplers)but instead were used once per location, in triplicate weeks.

Zoo species detection distances were estimated using Google Earth, calculating the mean detection distance per sampling method to the centroid of each species exhibit for each outdoor sampling location. Using these radii, we determined the percentage of ex-pected zoo species detected per sampling strategy over three weeks. Maximum detection distances were also averaged, omitting zeros to prevent overdispersion.

To examine the relationship between species biomass and detection probability (sensu [23,25]), we multiplied published mean biomass values (Supplementary Information, Table S2) by the number of individuals housed in the zoo. A Pearson correlation test was used to assess the relationship between total biomass and read counts.

## RESULTS

### Sequencing results

To detect airborne eDNA from vertebrates and mammals, we targeted mitochondrial regions using 12S and 16S markers. For the 12S dataset, we reduced our original 3,862,876 se-quencing reads to 783,731 reads across 293 ASVs after filtering, removing unidentifiable ASVs and correcting for negative controls. Subsequently, questionable species identifica-tions were adjusted using the weighted LCA algorithm in Galaxy. Despite our use of human blocking primers, 299,733 reads were identified as human (*Homo sapiens*), which we re-moved. This resulted in 483,998 reads across 161 ASVs, which we identified to 65 taxa,15 of which were expected resident species of the Rotterdam Zoo (total read count of 48,735 across 26 ASVs).

For 16S, we filtered 5,871,156 sequencing reads down to 651,690 reads after remov-ing unidentifiable ASVs and correcting for negative controls. Of the remaining reads, 293,267 were identified as *Homo sapiens*. After removal, the 16S dataset consisted of 358,249 reads across 92 ASVs that we identified to 47 taxa. Of these taxa, 23 were not identified using the 12S primer set, and 15 ASVs were identified as expected resident species of Rotterdam Zoo (total read count of 146,019 across 23 ASVs).

Cumulatively, we detected airborne eDNA from 88 distinct species, of which 24 were found using both primer sets (Supplementary Information, Table S3). Of these 24 species from both primer sets, six were expected zoo species: the Bactrian camel (*Camelus bactri-anus*), the black rhinoceros (*Diceros bicornis*), the giraffe (*Giraffa camelopardalis*), the Euro-pean otter (*Lutra lutra*), the Indian rhinoceros (*Rhinoceros unicornis*), and the Arctic fox (*Vulpes lagopus*).

### Species detection

We found that for overall species richness, sampling strategy (F_2_=66,87, p<0.001) but neither replicate week (replicates; F_2_=2.85, p=0.07) nor the interaction of sampling strategy across weeks (F_4_=0.54, p=0.70) was significantly different. Across all three sampling weeks (repli-cates) and samples acquired at the Rotterdam Zoo, we detected a total of 88 species: 34 mammals, 29 birds, 15 fish, and 10 amphibians (Figure 2). The passive Nutshell eDNA Sampler strategy accounted for 62.17% of all reads and detected 76 species, whereas the active sampler strategies, the Pollensniffer and Coriolis Micro, detected 50 species (21.66% of the reads) and 25 species (16.17% of reads), respectively. Of the species diversity de-tected, the majority (30) were exclusively collected by the Nutshell eDNA sampler strategy, and 17 were detected by all three approaches (Figure 2B). This pattern was similar when looking specifically at zoo residents (Figure 2C). Overall, 24 of the total 88 species detected using these air eDNA samplers were listed as Rotterdam Zoo residents: 15 mammals and 9 birds, accounting for 9% of the total inventory at Rotterdam Zoo (including those not on ex-hibit). Of these zoo species, the Nutshell eDNA sampler method detected 20 species, 9 of which were exclusive to this passive sampler (Figure 2C), whereas the Pollensniffer method collected 14 zoo species, of which 4 were exclusive to the Pollensniffer, and the Coriolis Mi-cro method detected 4 zoo species.

**Figure 2:**
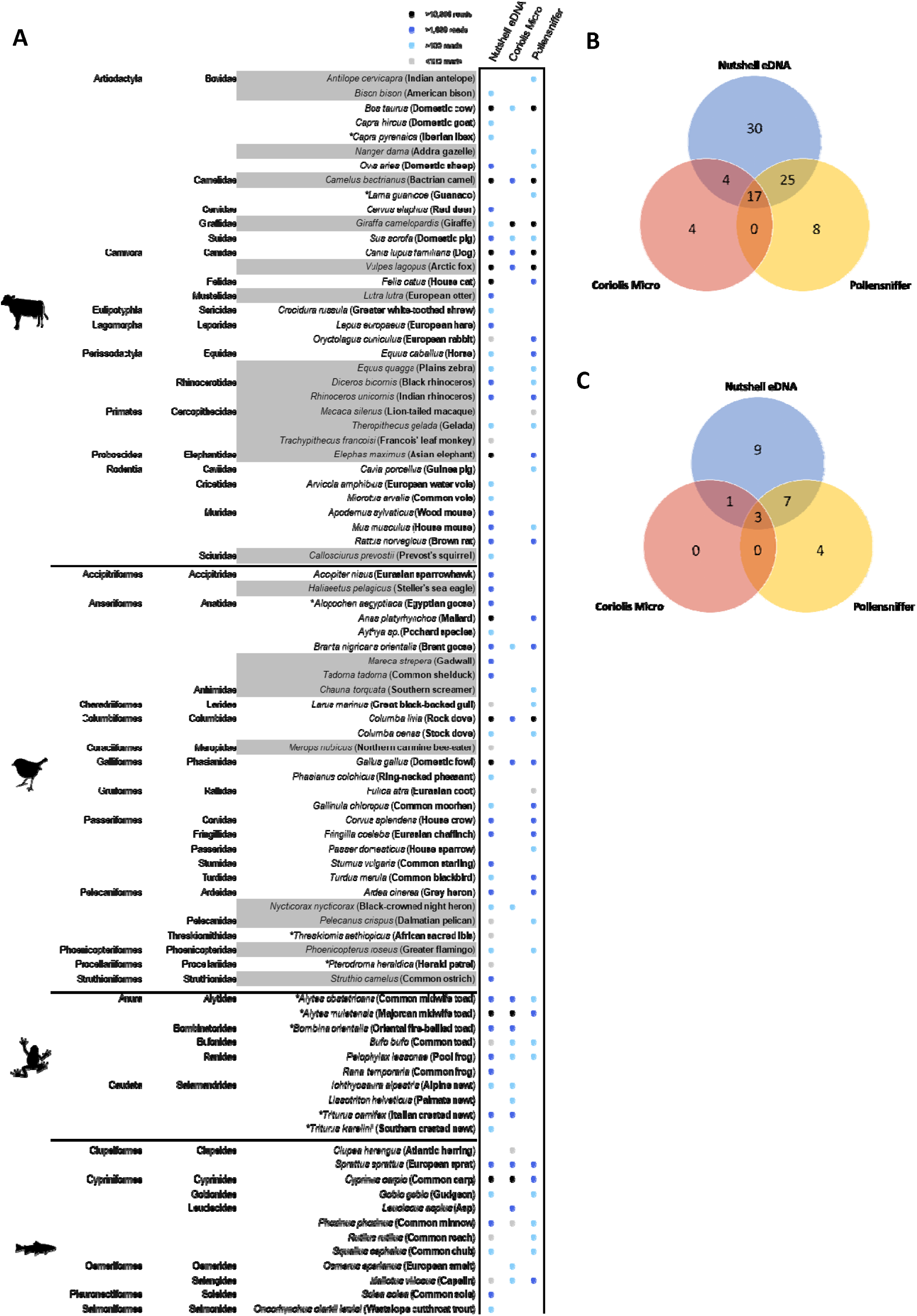
(**A)** Species diversity of taxa (icons per family) detected across three weeks of airborne eDNA sampling using different samplers: Nutshell eDNA sampler (passive sampler), Coriolis Micro, and Pollensniffer (active samplers). Total sequencing reads across all samples per sampler type indicated for each taxa by the corresponding circle colour (Legend top right).Grey rows represent resident zoo species. Asterisk denotes Netherlands non-native species**. (B)** Venn diagram of total species detection distribution across different air eDNA samplers**. (C)** Venn diagram of zoo resident species detection distribution across different air eDNA samplers.

While the majority of the non-target species (N = 64) were those native to the Nether-lands, including mammals, birds, amphibians and fish, 10 non-native species were also iden-tified (Figure 2). At the indoor sampling site, four species were detected that were not de-tected outdoors via any sampling strategy.

To accommodate the time series collections for our passive strategy (n = 90), the sampling intensity was not evenly comparable to each active strategy (n = 18). Thus, the per sample detection rate equated to 0.84 species/sample for the Nutshell eDNA sampler, 1.39 species/sample for the Coriolis Micro and 2.78 species/sampler for the Pollensniffer. When looking at zoo species specifically, the Nutshell eDNA sampler and Coriolis Micro each detected 0.22 zoo species/sample, whereas the Pollensniffer detected 0.78 zoo species/sample.

### Accumulating species richness detection over time

Comparing the three air eDNA sampler strategies, the mean total detected species richness by the Nutshell eDNA sampler per outdoor sampling location exceeded that of the Coriolis Micro (100L/min) between 6 and 24 hours (Figure 3), whereas the Pollensniffer at 8L/min was exceeded by the Nutshell eDNA sampler approach between 24 and 48 hours of passive sampling. The Nutshell eDNA sampler method detected on average 15.07 (SD ± 2.62) species per outdoor site after 96 hours of sampling (Figure 3SA), with site 2 detecting the most species diversity, on average 19 species (SD ± 5.29), and site 3 detecting the least number of species (11.67 SD ± 3.21), after 96 hours. Comparatively, the Pollensniffer detected an average of 6.6 species (SD ± 1.38), whereas the Coriolis Micro detected an average of 3.47 species (SD ± 1.15) per site after 10 minutes of active sampling. Excluding all non-target species, the mean total detected zoo species richness by the Nutshell eDNA sampler per outdoor sampling location exceeded that of the Coriolus Micro between 6 and 24 hours, whereas the Pollensniffer was exceeded by the Nutshell eDNA sampler approach between 48 and 72 hours of sampling (Figure 3B). We detected an average of 2.87 zoo species (SD ± 1.57) per outdoor sampling site after 96 hours of sampling using the Nutshell eDNA sampler (Figure S3B). By comparison, the Pollensniffer and the Coriolis Micro detected an average of 1.67 (SD ± 0.82) and 0.4 (SD ± 0.43) zoo species at each outdoor sampling site, respectively. The average detected zoo species richness by the Coriolis Micro was exceeded after 24 hours of sampling using the Nutshell eDNA sampler, and after approximately 72 hours using the Pollensniffer.

**Figure 3:**
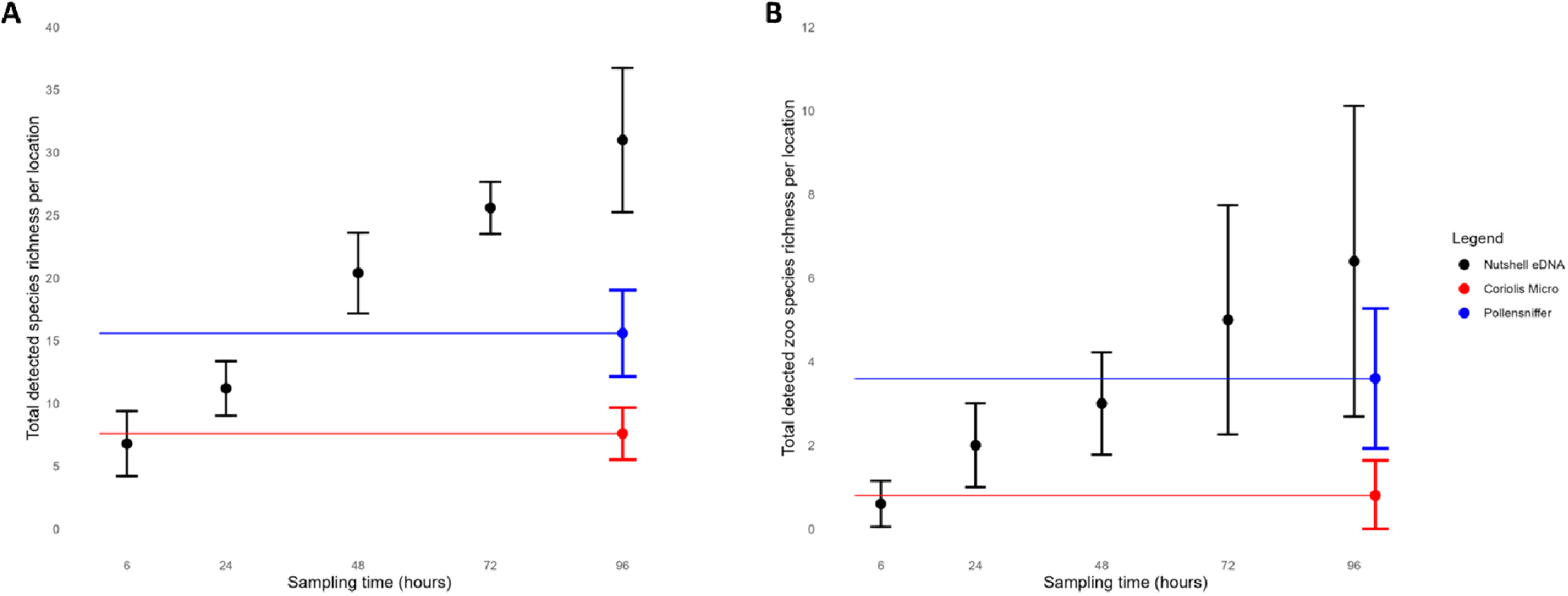
Species richness means (±SD) for outdoor sampling locations (sites 1-5). **(A)** Total detected species richness per outdoor sampling location by the Nutshell eDNA sampler strategy (black) over 96 hours across the three weeks of collection, juxtaposed against the mean (±SD) total detected species richness per outdoor sampling location by the Coriolis Micro (red) and Pollensniffer (blue) static samples per week in triplicate, with **(B)** showing total zoo resident species richness detection. Classified as Internal | Intern

Our indoor sampling site (site 6) demonstrated the mean detected species richness and zoo species richness obtained by the Coriolis Micro were both exceeded by the Nutshell eDNA sampler approach within the first 6 hours, similar to the outdoor sampling locations (Supplementary Information, Figure S4). In contrast, the Nutshell eDNA sampler strategy only exceeded the species richness obtained by the Pollensniffer method after 72 to 96 hours (Supplementary Information, Figure S4).

Accounting for triplicate measures across all three sampling strategies per location, our results indicate the Nutshell eDNA sampler approach detected significantly higher total species richness, followed by the Pollensniffer, and the Coriolis Micro detected the least (F(2,12) = 43.21, p < 0.001) (Supplementary Information, Figure S5A). The Coriolis Micro detected a total of 0.8 zoo species on average across three measurements per outdoor sampling location (SD ± 0.84), whereas the Pollensniffer approach detected a total of 3.6 (SD ± 1.67) zoo species on average across three measurements per outdoor sampling location (Figure S3B).

Our results demonstrate the total detected zoo species per outdoor sampling location to be significantly different among all three strategies (F(2,12) = 6.80, p = 0.011), with the highest total zoo species richness detected by the Nutshell eDNA sampler (Supplementary Information, Figure S5B).

Equalizing sampling intensity, we also individually analysed each collection duration time that the Nutshell eDNA sampler was deployed (6, 24, 28, 72 and 96 hours) and compared it to the species richness obtained from both active approaches (Figure. 4A). Although total species richness detected across all outdoor sites and replicates across time was 74 species for the Nutshell eDNA sampler, 50 species for the Pollensniffer and 25 species for the Coriolis Micro, we observed the Nutshell eDNA sampler to detect fewer species than the Pollensniffer at each independent collection time point when standardizing sampler number (n = 3 per location, per deployment time). While the Nutshell eDNA sampler accumulated species richness over time across the outdoor locations, from 19 species when deployed for 6 hours to 38 species for 96 hours (Figure 4A), only 23% of total species richness detected (n = 17) was retained continuously from the time of first detection until the end of sampling at 96 hours (Figure 4B). Interestingly, while 12.2% (n = 9) of identified taxa were detected across every time point taken for the Nutshell eDNA sampling strategy, 51.4% (n = 38) were only identified during a single time point (“singletons”) and another 25.7% (n = 19) intermittently or “patchy” detections (>2 time points).

**Figure 4:**
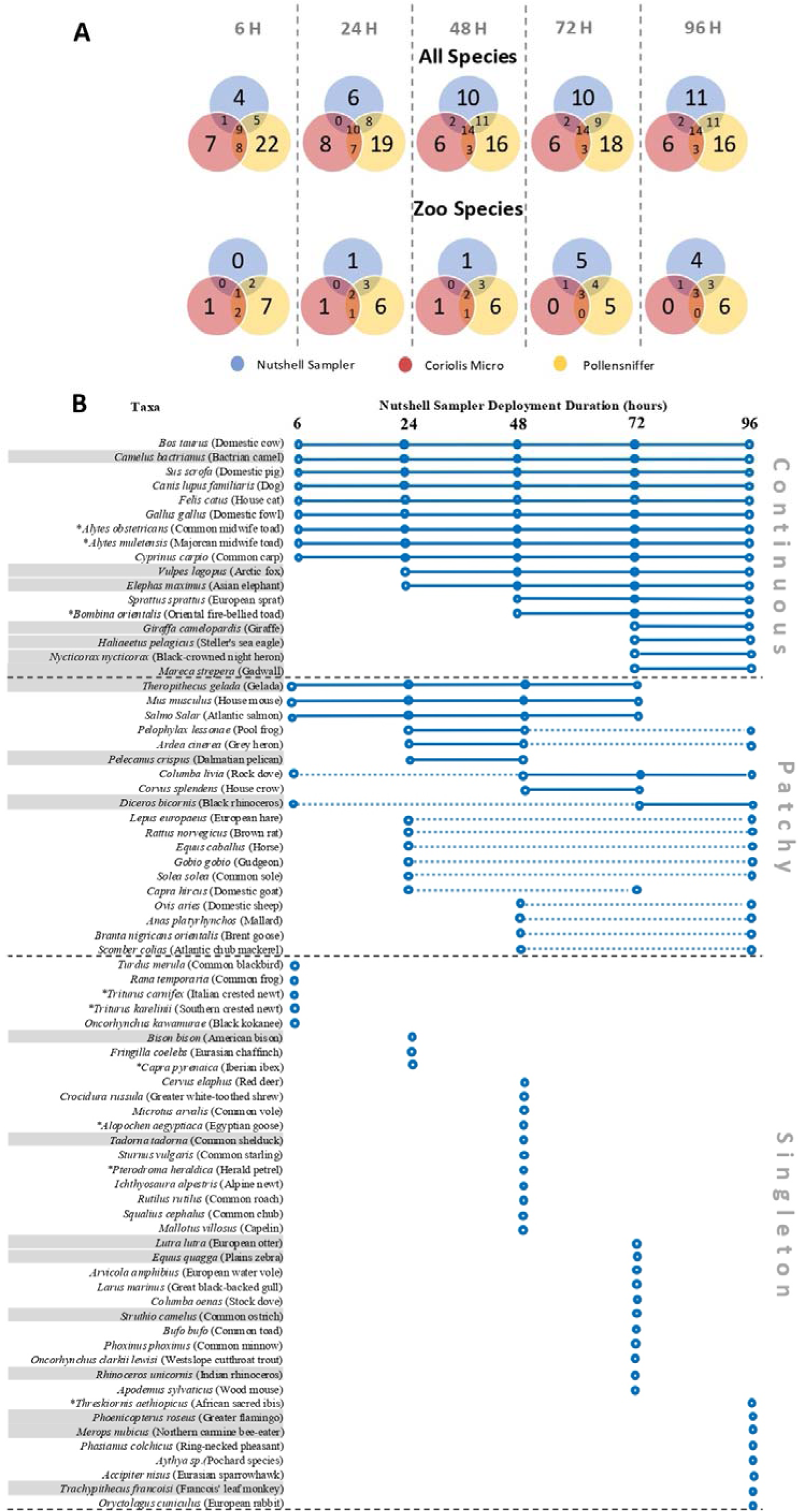
(A) Venn diagrams of species richness detected by different air eDNA strategies (bottom legend) across time (6, 24, 48, 72, 96 hours) for all species detected (above) and zoo resident species only (bottom). Note that active air eDNA samples were not collected at each timepoint; the apparent switching in the number of unique detections between timepoints reflects passive samplers intermittently capturing additional species that overlap with those detected by the active samplers. **(B)** Species-specific detection over time (6, 24, 48, 72, 96 hours) across outdoor sites for passive Nutshell eDNA sampler. Blue circles denote detection at time point, solid lines signify continuous detections (“continuous”), dashed lines denote no detections between positive detections (“patchy”), compared to single detections (“singletons”). Grey rows represent resident zoo species. Asterisk denotes Netherlands non-native species.

Similar patterns were mirrored for resident zoo species (n=19), wherein 58% of total species richness was detected at the 96-hour deployment time for the Nutshell eDNA sampler (Figure 4A). Of the 19 resident zoo species detected by the Nutshell eDNA sampler, 36.8% (n=7) of species signal were retained for the entire deployment time (96 hours), with 15.8% (n=3) presenting “patchy” signals, and 47.4% (n=9) representing “singleton” detections (Figure 4B).

### Accuracy, dispersion, and biomass

The Nutshell eDNA sampler detected zoo species at the greatest average distance (188.67 ± 61.33 m), followed by the Pollensniffer (172.58 ± 91.29 m), and the Coriolis Micro (68 ± 50.21 m), though these differences were not statistically significant (F(2,10) = 2.8, *p* = 0.11) (Figure 5A; Supplementary Information, Table S4). Excluding the indoor site, the Nutshell eDNA sampler also recorded the furthest detections (373 ± 124.37 m) with a maximum detection of an American bison (*Bison bison*) at 515 m (location 2), which were significantly higher than the furthest detections of the Coriolis Micro (*p* = 0.036), (Figure 5B). The furthest detections by the Pollensniffer and Coriolis Micro were an Arctic fox (*Vulpes lagopus*, 491 m) and a black-crowned night heron (*Nycticorax nycticorax*, 128 m) respectively, however both at the indoor location (location 6). Outdoors, the farthest detections were the Asian elephant (*Elephas maximus*, 466 m, Pollensniffer) and the night heron again (116 m, Coriolis Micro).

**Figure 5:**
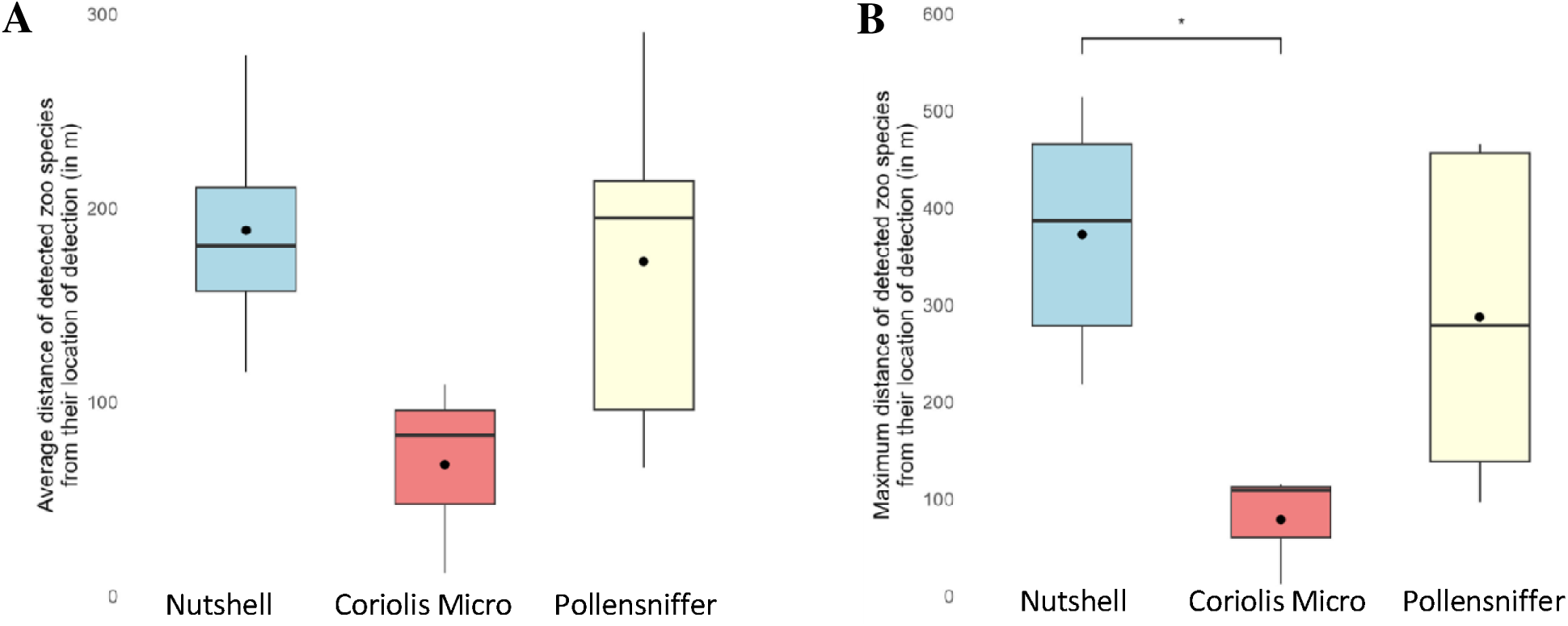
Boxplots showing the average (A) and maximum (B) distance in meters between the detected zoo species and the sampling location at which they were detected for the different outdoor sampling locations per air eDNA sampler across three weeks of sampling. Asterix denotes significant different at p<0.05.

In terms of detection accuracy, the Nutshell eDNA sampler strategy demonstrated the highest results, identifying 19% (±11.5%) of zoo species within its average range, followed by the Pollensniffer (12% ± 5.8%) and the Coriolis Micro (16,6% ± 23,5%). Using the lowest average detection distance (68 m, Coriolis Micro) as a baseline, both the Nutshell eDNA sam-pler method (33.2% ± 20.4%) and the Pollensniffer approach (21.6% ± 21.7%) showed higher detection accuracy across five outdoor locations, indicating effective identification of nearby zoo species (Figure 6). Additionally, there was a significant positive correlation be-tween total sequencing reads from our air eDNA samples and the biomass (log-transformed) of detected zoo species (r(24) = 0.44, *p* = 0.031; Supplementary Information Fig. S5a). This trend remained, though weaker, after removing four exceptionally large outliers (r(20) = 0.38, *p* = 0.097; Supplementary Information, Fig. S6B).

**Figure 6:**
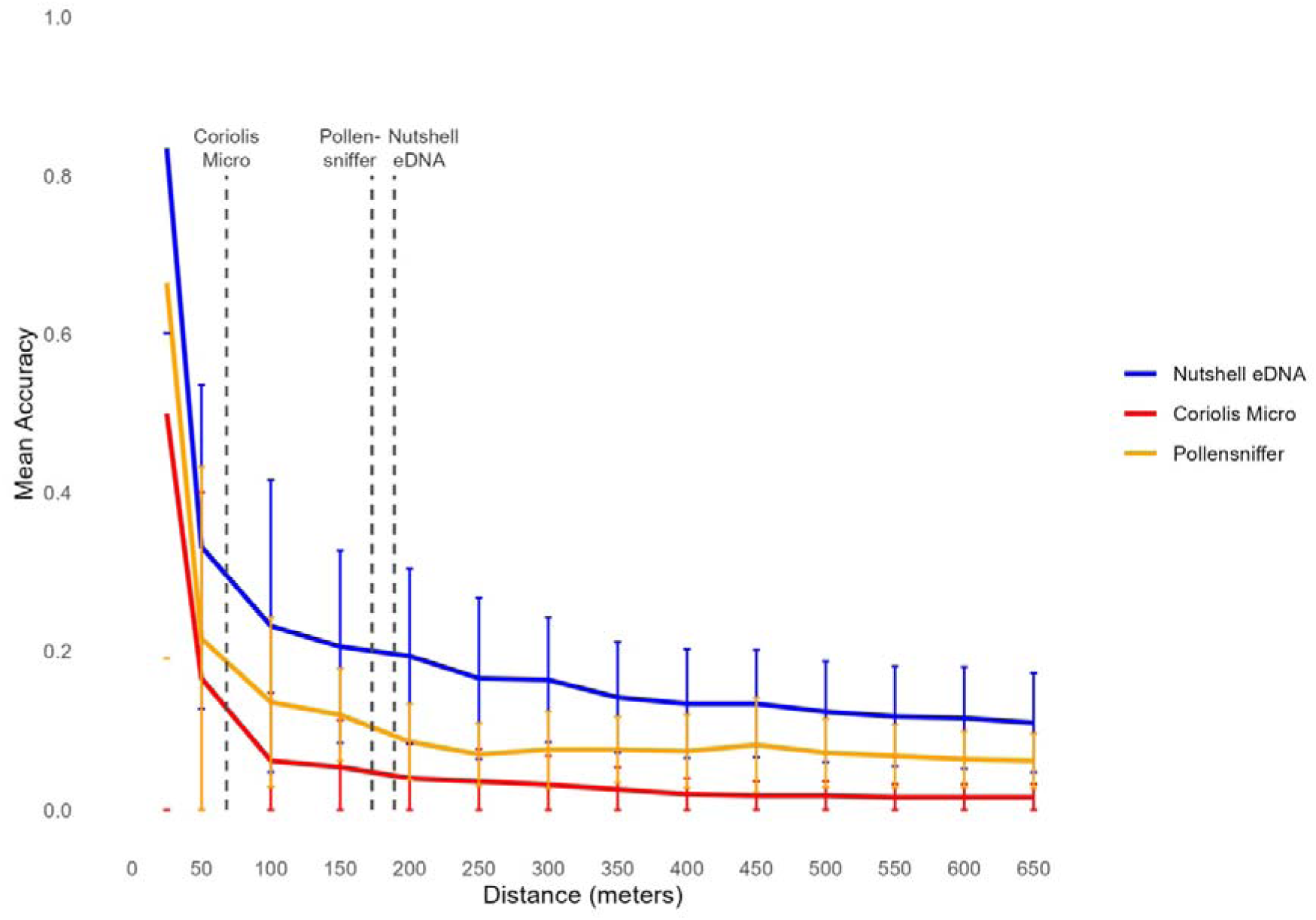
Mean % detection accuracy dispersion curve across distance (meters ± SD) from sampling location of detected zoo species. Dashed lines represent each sampler’s average distance (meters). Sampler type is denoted by colour (legend to the right).

## DISCUSSION

Our study aimed to assess airborne eDNA sampling strategies by comparing the effi-ciency of a newly designed, 3D-printed, passive approach and two commonly used active methods, while exploring the spatial and temporal dynamics of airborne eDNA. Our results show that the passive strategy detected a broad range of zoo and non-zoo species, and in total captured a higher variety of vertebrate species over a 96-hour integrated sampling pe-riod than either active sampler strategy could given their time-point measurement approach. Specifically, while the active sampling strategy employing the Pollensniffer (10 minutes of sampling) detected the most species richness per sampling effort, the passive strategy using the Nutshell eDNA sampler accumulated species richness over deployment time while also detecting unique species at different time points. Although mean detection distances did not differ significantly, the Nutshell eDNA sampler method identified taxa at the furthest range and achieved the highest accuracy within its average detection radius. Additionally, we found a positive correlation between species biomass and total sequencing reads obtained from airborne eDNA, suggesting a scaling relationship to eDNA released into the air such that large vertebrates (or their populations) release substantially more eDNA into the air, puta-tively offering insights into vertebrate relative abundance.

In total, this investigation suggests our passive air eDNA method may offer an alter-native or a complimentary method to active air DNA sampling strategies when hung for as little as a few days, also enabling the detection of intermittent or patchy air eDNA signals that may not be detected through static, short-burst sampling approaches like that of active sampling.

### Species detection within Rotterdam Zoo

We detected airborne eDNA from 88 vertebrate species, spanning both terrestrial (mammals, birds) and aquatic (fish, amphibians) habitats. Of these, 76 species were detected by the passive Nutshell eDNA sampler within 96 hours, including 30 not detected by either active sampler highlighting effective detection capacity of passive sampling. While the Pollensniffer showed similar diversity to previous active air eDNA studies in zoos [23,25], 64 of the 88 taxa were non-target species, likely local wildlife or food (e.g., livestock, fish).

The question remains as to why our passive airborne eDNA sampler detected a wider array of species compared to both active samplers. When accounting for per sample detection rates, the Pollensniffer detected the highest recovery of species richness but the other active sampler, the Coriolis Micro, detected the least. While in part this implies more sampling effort alone increases the likelihood for increasing species richness detections, when sampling intensity was normalized, the passive strategy outperformed at least one of the two active strategies. Currently, little is known about the movement of airborne eDNA, and the impact of sampling strategies. While some species were detected across multiple samples and sampling sites (e.g. *Cyprinus caprio* in 58 samples; *Elephas maximus* in 28 samples), 32 species were present in only a single sample, highlighting the likely stochastic nature in detecting airborne eDNA in general. The greater detection capacity of the Nutshell eDNA sampler may stem from factors like sampling height (∼2.5 m vs. ≤1 m for active samplers), extended sampling duration (96 hours vs. short active bursts), and despite our inability to quantify such a metric at present, a likely higher cumulative air volume sampled. This longer, passive sampling approach may integrate effects tied to eDNA release variability due to species behaviour or environmental factors (e.g., wind, rain), aggregating potential eDNA release bursts and thus increasing species detection over other methods [51].

At the indoor site (location 6), however, active sampling by the Pollensniffer outper-formed our passive method possibly due to the limited airflow indoors which hampers pas-sive collection but promotes airborne eDNA accumulation [7, 52]. In contrast but similar to the outdoor sites, the Coriolis Micro detected the lowest diversity despite its higher flow rate. Notably, the Pollensniffer consistently detected more species per location than the Coriolis Micro (Fig. S3a), despite sampling less air volume per minute. This may reflect differences in sampling mechanics; the impingement method of the Pollensniffer might better capture fine aerosols than the vortex mechanism of the Coriolis Micro, where high airflow could reduce DNA particle retention. Further studies using matched air volumes but varying flow rates are needed to test this hypothesis [53].

### Accumulation patterns of airborne eDNA

Per site, the Nutshell eDNA sampler outperformed the Coriolis Micro in species richness within the first 24 hours, both overall and for zoo species. Compared to the Pollensniffer, it surpassed species richness after 24–48 hours (overall) and 48–72 hours (zoo species).

Given that many active samplers are noisy, labour-intensive, costly, and might require tech-nically trained personnel, passive sampling offers a low-cost, minimally intrusive alternative.

Consistent with [30] who showed airborne eDNA accumulates over 24 hours using water-based passive samplers, we observed no plateau in species detection even after 96 hours. Continuous detection of new species suggests that longer sampling durations could further increase species richness. Unlike aquatic eDNA, which may be more homogenous, airborne eDNA might be released in pulses or be influenced by weather, as suggested by [28] and [53]. Indeed, we observed variation in both species richness and composition over time, highlighting the potential impact of pulsed release, weather, and eDNA persistence on airborne eDNA detection signatures. While many species were detected from the outset of our deployment time (6 hours) and retained until the last deployment time (96 hours), many species also demonstrated “patchy” detections wherein focal species were detected at two or more time points but not retained for the entire deployment time. Similarly, many species represented “singleton” detections that were also not captured in samplers hung for other durations. This may suggest either (i) a potential trade-off between species accumulation and retention on filters, (ii) species-specific DNA signals are emitted at specific time intervals such that the sampler detected pulsed or intermittent DNA releases, or (iii) air eDNA is not homogeneously mixed and thus paired samplers at the same site did not collect the same air eDNA particulate matter despite it being potentially available.

A trade-off between DNA signal accumulation over time and lost DNA signal retention due to degradation is certainly likely given our general knowledge of the ecology of eDNA, at least within aquatic systems [54, 55]. However, given some of our detected species (23%, total; 36.8% zoo) did retain their signal until 96 hours of deployment time (Figure 4B) and the known high efficiency of glass fibre filters in bioaerosol collections [56], we suspect these patterns to be more indicative of either the capture of biologically intermittent signals, or al-ternatively, the heterogenous distribution of airborne eDNA within the airstream. Untangling these effects experimentally and devising optimized sampling strategies will be important for future studies, particularly for differentiating accumulation, persistence and retention of signals after their collection.

### Spatial distribution of target species

Out of all airborne eDNA sampling strategies, the passive approach had the highest average detection distance and accuracy, identifying nearly 1 in 5 target species within its 188m range. Both the Nutshell eDNA sampler and Pollensniffer were more effective at detecting nearby species than the Coriolis Micro, aligning with findings from [23] and [25], who observed stronger detection closer to source animals. While airborne eDNA typically concentrates around its origin, it can also travel considerable distances, such as the American bison (*Bison bison*) detected 515m from its enclosure by the Nutshell eDNA sampler within 24 hours. Notably, the furthest detections by the active samplers occurred indoors, underscoring the challenge of tracing eDNA back to its source in natural settings [7] and the high potential for allochthonous signal detection. Similar difficulties have occasionally been reported in aquatic systems for localizing sources of eDNA (e.g., [57]), highlighting challenges regardless of the environmental media sampled. One possible explanation is that indoor environments act as sinks for airborne eDNA. As doors open and close, and as people, including visitors and researchers, move through these spaces, they may introduce and resuspend previously settled eDNA. Air circulation systems such as air conditioners or ceiling fans may further aerosolize particles, as discussed by [7] who documented complex airborne eDNA signatures in indoor tropical environments. These dynamics suggest that eDNA can readily enter indoor spaces but may persist longer due to limited dispersal mechanisms, with implications for both spatial resolution and temporal acuity. In contrast, outdoor eDNA may be more prone to dilution or rapid dissipation, potentially reducing signal persistence but improving spatial specificity.

Conversely, many nearby target species went undetected, with contributing factors likely being particle heterogeneity, wind patterns, UV degradation, low species-specific shedding rates, or a lack of reference sequences [7, 58]. Echoing findings from aquatic studies 59], zoo-based airborne eDNA research has shown a correlation between species and detection likelihood [23,25]. Our results similarly revealed a positive size-scaling relationship between air eDNA abundance and the total species biomass housed in populations at the zoo, suggesting larger vertebrates might be more likely to be detected with air eDNA, a feature that is becoming a salient characteristic for both aquatic and air eDNA.

### Future directions and recommendations

This study is the first to apply the passive Nutshell eDNA sampler for airborne eDNA-based vertebrate detection, though its optimal setup remains to be determined. We used 0.7 µm glass fibre filters, but other materials may enhance detection. For instance, [25] used F8 pleated filters, and [23] used Sterivex-HV filters (0.22–0.45 µm), finding that pore size did not significantly affect community composition. Since the Nutshell eDNA sampler passively col-lects particles in a glass fibre matrix without forcing air through pores, pore size may be less critical than with active samplers.

To date, several variables remain unexplored: filtering duration, seasonal effects, sampler height, and air movement. Wind direction is thought to influence detection more dur-ing passive sampling [60], though we did not track it in this study. Notwithstanding the above, the Nutshell eDNA sampler effectively detected both zoo and local species and continued to register new detections over time suggesting optimizing for technical and biological influ-ences are likely to only improve detection and biodiversity resolution.

## CONCLUSION

Overall, the passive strategy for collecting air eDNA offers a low-cost, non-invasive, and effi-cient method for vertebrate biodiversity assessment when compared to its active strategy counterparts. Its ease of deployment, reliance on airstream absorption rather than particle settlement, and wide taxonomic detection range position it a valuable tool, especially for ex-panding terrestrial biomonitoring. Future research should refine passive sampling protocols and evaluate their performance in natural habitats. While terrestrial eDNA research currently lags behind aquatic systems, airborne eDNA has transformative potential for large-scale bio-diversity monitoring. Given its strong detection performance across taxa, passive air eDNA is an underappreciated yet powerful method for biodiversity assessments and may emerge as the preferred strategy for future terrestrial biomonitoring, particularly for scaling-up efforts (e.g., via citizen science).

## ACKNOWLEDGEMENTS

This research was made posible through the NWO Aspasia funding (KAS) and research access granted through Blijdorp Zoo.

## DATA ARCHIVING

All data can be found at FigShare DOI: 10.6084/m9.figshare.28842980

## REFERENCES

1. Tong, S.; Bambrick, H.; Beggs, P. J.; Chen, L.; Hu, Y.; Ma, W.; Steffen, W.; Tan, J. Current and Future Threats to Human Health in the Anthropocene. Environ. Int. 2022, 158, 106892. 10.1016/j.envint.2021.106892.

2. Zimmerer, K. S.; de Haan, S.; Jones, A. D.; Creed-Kanashiro, H.; Tello, M.; Ca-rrasco, M.; Meza, K.; Plasencia Amaya, F.; Cruz-Garcia, G. S.; Tubbeh, R.;, et al. The Biodiversity of Food and Agriculture (Agrobiodiversity) in the Anthropocene: Research Advances and Conceptual Framework. Anthropocene 2019, 25, 100192. 10.1016/j.ancene.2019.100192.

3. Baird, D. J.; Hajibabaei, M. Biomonitoring 2.0: A New Paradigm in Ecosystem As-sessment Made Possible by Next-Generation DNA Sequencing. Mol. Ecol. 2012, 21(8), 2039–2044. 10.1111/j.1365-294X.2012.05519.x.

4. Rodríguez-Ezpeleta, N., Zinger, L., Kinziger, A., Bik, H. M., Bonin, A., Coissac, E., Emerson, B. C., Lopes, C. M., Pelletier, T. A., Taberlet, P., & Narum, S. (2021). Biodiversity monitoring using environmental DNA. Molecular Ecology Resources, 21(5), 1405–1409. 10.1111/1755-0998.13399

5. Taberlet, P.; Coissac, E.; Pompanon, F.; Brochmann, C.; Willerslev, E. Towards Next-Generation Biodiversity Assessment Using DNA Metabarcoding. Mol. Ecol. 2012, 21(8), 2045–2050. 10.1111/j.1365-294X.2012.05470.x.

6. Didaskalou, E. A.; Trimbos, K. B.; Stewart, K. A. Environmental DNA. Curr. Biol. 2022, 32(22), R1250–R1252. 10.1016/j.cub.2022.09.052.

7. Garrett, N. R., Watkins, J., Francis, C. M., Simmons, N. B., Ivanova, N., Naaum, A., Briscoe, A., Drinkwater, R., & Clare, E. L. (2023). Out of thin air: surveying tropical bat roosts through air sampling of eDNA. PeerJ, 11, e14772. 10.7717/peerj.14772

8. Kelly, R. P., Port, J. A., Yamahara, K. M., Martone, R. G., Lowell, N., Thomsen, P. F., Mach, M. E., Bennett, M., Prahler, E., Caldwell, M. R., & Crowder, L. B. (2014). Harnessing DNA to improve environmental management. Science, 344(6191), 1455–1456. 10.1126/science.1251156

9. Qu, C., & Stewart, K. A. (2019). Evaluating monitoring options for conservation: comparing traditional and environmental DNA tools for a critically endangered mammal. The Science of Nature, 106(3–4), 9. 10.1007/s00114-019-1605-1

10. Hallam, J., Clare, E. L., Jones, J. I., & Day, J. J. (2021). Biodiversity assessment across a dynamic riverine system: A comparison of eDNA metabarcoding versus traditional fish surveying methods. Environmental DNA, 3(6), 1247–1266. 10.1002/edn3.241

11. Fediajeviate, J., Priestly, V., Arnold, R., Savolainen, V. 2021. Meta-analysis shows that environmental DNA outperforms traditional surveys but warrants better re-porting standards. Ecology and Evolution. 11(9), 4803–4815. 10.1002/ece3.7382

12. Ruppert, K. M., Kline, R. J., & Rahman, M. S. Past, present, and future perspec-tives of environmental DNA (eDNA) metabarcoding: A systematic review in methods, monitoring, and applications of global eDNA. Global Ecology and Conservation (2019), 17, e00547. 10.1016/j.gecco.2019.e00547

13. Gold, Z., Sprague, J., Kushner, D. J., Zerecero Marin, E., & Barber, P. H. (2021). eDNA metabarcoding as a biomonitoring tool for marine protected areas. PLOS ONE, 16(2), e0238557. 10.1371/journal.pone.0238557

14. Thomsen, P. F.; Kielgast, J.; Iversen, L. L.; Møller, P. R.; Rasmussen, M.; Willerslev, E. Detection of a Diverse Marine Fish Fauna Using Environmental DNA from Seawater Samples. PLoS ONE 2012, 7(8), e41732. 10.1371/journal.pone.0041732.

15. van Kuijk, J.L., van den Burg, M.P., Didaskalou, E.A., et al. Safeguarding *Iguana* diversity: enabling rapid and low-effort tracking of non-native iguanas through terrestrial eDNA innovations. Biol Invasions 27, 63 (2025). 10.1007/s10530-024-03524-x

16. Beng, K. C.; Corlett, R. T. Applications of Environmental DNA (eDNA) in Ecology and Conservation: Opportunities, Challenges, and Prospects. Biodivers. Conserv. 2020, 29, 2089–2121. 10.1007/s10531-020-01980-0.

17. Kinoshita, G., Yonezawa, S., Murakami, S., & Isagi, Y. (2019). Environmental DNA Collected from Snow Tracks is Useful for Identification of Mammalian Species. Zoological Science, 36(3), 198. 10.2108/zs180172

18. Lyman, J. A., Sanchez, D. E., Hershauer, S. N., Sobek, C. J., Chambers, C. L., Zahratka, J., & Walker, F. M. (2022). Mammalian eDNA on herbaceous vegetation? Validating a qPCR assay for detection of an endangered rodent. Environmental DNA, 4(5), 1187–1197. 10.1002/edn3.331

19. Thomsen, P. F.; Sigsgaard, E. E. Environmental DNA Metabarcoding of Wild Flowers Reveals Diverse Communities of Terrestrial Arthropods. Ecol. Evol. 2019, 9(4), 1665–1679. 10.1002/ece3.4809.

20. Sigsgaard, E. E., Olsen, K., Hansen, M. D. D., Hansen, O. L. P., Høye, T. T., Svenning, J., & Thomsen, P. F. Environmental DNA metabarcoding of cow dung reveals taxonomic and functional diversity of invertebrate assemblages. Molecular Ecology (2021), 30(13), 3374–3389. 10.1111/mec.15734

21. Roger, F., Ghanavi, H. R., Danielsson, N., Wahlberg, N., Löndahl, J., Pettersson, L. B., Andersson, G. K. S., Boke Olén, N., & Clough, Y. (2022). Airborne environmental DNA metabarcoding for the monitoring of terrestrial insects—A proof of concept from the field. Environmental DNA 2014, 4(4), 790–807. 10.1002/edn3.290

22. Clare, E. L., Economou, C. K., Faulkes, C. G., Gilbert, J. D., Bennett, F., Drinkwater, R., & Littlefair, J. E. (2021). eDNAir: proof of concept that animal DNA can be collected from air sampling. PeerJ, 9, e11030. 10.7717/peerj.11030

23. Clare, E. L., Economou, C. K., Bennett, F. J., Dyer, C. E., Adams, K., McRobie, B., Drinkwater, R., & Littlefair, J. E. (2022). Measuring biodiversity from DNA in the air. Current Biology, 32(3), 693–700.e5. 10.1016/j.cub.2021.11.064

24. Gusareva, E. S., Acerbi, E., Lau, K. J. X., Luhung, I., Premkrishnan, B. N. V., Kolundžija, S., Purbojati, R. W., Wong, A., Houghton, J. N. I., Miller, D., Gaultier, N. E., Heinle, C. E., Clare, M. E., Vettath, V. K., Kee, C., Lim, S. B. Y., Chénard, C., Phung, W. J., Kushwaha, K. K., … Schuster, S. C. (2019). Microbial communities in the tropical air ecosystem follow a precise diel cycle. Proceedings of the National Academy of Sciences of the United States of America, 116(46), 23299–23308. 10.1073/pnas.1908493116

25. Lynggaard, C., Bertelsen, M. F., Jensen, C. V., Johnson, M. S., Frøslev, T. G., Olsen, M. T., & Bohmann, K. (2022). Airborne environmental DNA for terrestrial ver-tebrate community monitoring. Current Biology, 32(3), 701–707.e5. 10.1016/j.cub.2021.12.014

26. Polling, M., Buij, R., Laros, I., & de Groot, A. (2024). Continuous daily sampling of airborne eDNA detects all vertebrate species identified by camera traps. Environmental DNA, 6(4), Article e591. 10.1002/edn3.591

27. Garrett, N. R., Watkins, J., Simmons, N. B., Fenton, B., Maeda-Obregon, A., Sanchez, D. E., Froehlich, E. M., Walker, F. M., Littlefair, J. E., & Clare, E. L. (2022). Airborne eDNA documents a diverse and ecologically complex tropical bat and other mammal community. Environmental DNA, 5(2), 350–362. 10.1002/edn3.385

28. Johnson, M. D.; Barnes, M. A.; Garrett, N. R.; Clare, E. L. Answers Blowing in the Wind: Detection of Birds, Mammals, and Amphibians with Airborne Environmental DNA in a Natural Environment Over a Yearlong Survey. Environ. DNA (2023), 5(2), 375–387. 10.1002/edn3.388.

29. Johnson, M. D., Fokar, M., Cox, R. D., & Barnes, M. A. (2021). Airborne environmental DNA metabarcoding detects more diversity, with less sampling effort, than a traditional plant community survey. BMC Ecology and Evolution, 21(1), 218. 10.1186/s12862-021-01947-x

30. Klepke, M. J., Sigsgaard, E. E., Jensen, M. R., Olsen, K., & Thomsen, P. F. (2022). Accumulation and diversity of airborne, eukaryotic environmental DNA. Environmental DNA, 4(6), 1323–1339. 10.1002/edn3.340

31. Newton, J. P., Nevill, P., Bateman, P. W., Campbell, M. A. & Allentoft, M. E. Spider webs capture environmental DNA from terrestrial vertebrates. iScience 27, 108904 (2024).

32. Bessey, C., Neil Jarman, S., Simpson, T. et al. Passive eDNA collection enhances aquatic biodiversity analysis. Commun Biol 4, 236 (2021). 10.1038/s42003-021-01760-8

33. Cananzi G, Tatini I, Li T, Montagna M, Serra V, Petroni G. Active or passive? A multi-marker approach to compare active and passive eDNA sampling in riverine environments. Sci Total Environ. 2025 Apr 25;974:179247. doi: 10.1016/j.scitotenv.2025.179247.

34. Lynggaard, C., Guldberg Frøslev, T., Johnson, M.S., Tange Olsen, M., Bohmann K. 2023. Airborne environmental DNA captures terrestrial vertebrate diversity in nature. Molecular Ecology Resources 24(1). E13840. 10.1111/1755-0998.13840

35. Johnson M, Barnes MA. (2023). Macrobial airborne environmental DNA analysis: A review of progress, challenges, and recommendations for an emerging application. Mol Ecol Resour. 2024 Oct;24(7):e13998. doi: 10.1111/1755-0998.13998.

36. de Weger, L. A.; Molster, F.; de Raat, K.; den Haan, J.; Romein, J.; van Leeuwen, W.; de Groot, H.; Mostert, M.; Hiemstra, P. S. A New Portable Sampler to Monitor Pollen at Street Level in the Environment of Patients. Sci. Total Environ. 2020, 741, 140404. 10.1016/j.scitotenv.2020.140404.

37. Bertin Technologies SAS. Coriolis Micro – Microbial Air Sampler. Available at: https://www.bertin-technologies.com/product/air-samplers/coriolis-micro-air-sampler/ (accessed March 2, 2025).

38. Luursema, J. M. (2023). Nutshell-eDNA-sampler. Available at: https://github.com/J4n-M44rt3n/Nutshell-eDNA-sampler [accessed March 2, 2025].

39. Yates, M. C.; Glaser, D. M.; Post, J. R.; Cristescu, M. E.; Fraser, D. J.; Derry, A. M. The Relationship between eDNA Particle Concentration and Organism Abundance in Nature is Strengthened by Allometric Scaling. Mol. Ecol. 2021, 30(13), 3068–3082. 10.1111/mec.15543.

40. Taylor, P. G. Reproducibility of Ancient DNA Sequences from Extinct Pleistocene Fauna. Mol. Biol. Evol. 1996, 13(1), 283–285. 10.1093/oxfordjournals.molbev.a025566.

41. Riaz, T., Shehzad, W., Viari, A., Pompanon, F., Taberlet, P., & Coissac, E. (2011). ecoPrimers: inference of new DNA barcode markers from whole genome sequence analysis. Nucleic Acids Research, 39(21), e145–e145. 10.1093/nar/gkr732

42. Vestheim, H.; Jarman, S. N. Blocking Primers to Enhance PCR Amplification of Rare Sequences in Mixed Samples – A Case Study on Prey DNA in Antarctic Krill Stomachs. Front. Zool. 2008, 5(1), 12. 10.1186/1742-9994-5-12.

43. Calvignac-Spencer, S., Merkel, K., Kutzner, N., Kühl, H., Boesch, C., Kappeler, P. M., Metzger, S., Schubert, G., & Leendertz, F. H. (2013). Carrion fly-derived DNA as a tool for comprehensive and cost-effective assessment of mammalian biodiversity. Molecular Ecology, 22(4), 915–924. 10.1111/mec.12183

44. Martin, M. (2011). Cutadapt removes adapter sequences from high-throughput sequencing reads. EMBnet.Journal, 17(1), 10. 10.14806/ej.17.1.200

45. Callahan, B. J.; McMurdie, P. J.; Rosen, M. J.; Han, A. W.; Johnson, A. J. A.; Holmes, S. P. DADA2: High-Resolution Sample Inference from Illumina Amplicon Data. Nat. Methods 2016, 13(7), 581–583. 10.1038/nmeth.3869.

46. RStudio Team (2020). RStudio: integrated development for R. http://www.rstudio.com/.

47. Afgan, E.; Baker, D.; Batut, B.; van den Beek, M.; Bouvier, D.; Čech, M.; Chilton, J.; Clements, D.; Coraor, N.; Grüning, B. A.; Guerler, A.; Hillman-Jackson, J.; Hiltemann, S.; Jalili, V.; Rasche, H.; Soranzo, N.; Goecks, J.; Taylor, J.; Nekrutenko, A.; Blankenberg, D. The Galaxy Platform for Accessible, Reproducible, and Collaborative Biomedical Analyses: 2018 Update. Nucleic Acids Res. 2018, 46(W1), W537–W544. 10.1093/nar/gky379.

48. Beentjes, K. K.; Barmentlo, S. H.; Cieraad, E.; Schilthuizen, M.; van der Hoorn, B. B.; Speksnijder, A. G. C. L.;, et al. Environmental DNA Metabarcoding Reveals Comparable Responses to Agricultural Stressors on Different Trophic Levels of a Freshwater Community. Mol. Ecol. 2021, 31, 1430–1443. 10.1111/MEC.16326.

49. Oksanen, A.J., Blanchet, F.G., Friendly, M., Kindt, R., Legendre, P., Mcglin, D., Minchin, P.R., Hara, R. B. O., Simpson, G.L., Solymos, P., Stevens, M. H. H., & Szoecs, E. (2016). Package ‘vegan’ January.

50. Wickham, H. ggplot2: Elegant Graphics for Data Analysis; Springer-Verlag: New York, 2016. https://ggplot2.tidyverse.org.

51. Lin, R., Yan, Z. & Yao, M. Robust passive sampling of airborne environmental DNA to monitor plants and animals. Methods Ecol. Evol. 16, 2118–2130 (2025).

52. Serrao, N. R., Weckworth, J. K., McKelvey, K. S., Dysthe, J. C., & Schwartz, M. K. Molecular genetic analysis of air, water, and soil to detect big brown bats in North America. Biological Conservation (2021), 261, 109252. 10.1016/j.biocon.2021.109252

53. Manibusan, S. & Mainelis, G. Passive bioaerosol samplers: A complementary tool for bioaerosol research. A review. J. Aerosol Sci. 163, 105992 (2022).

54. Barnes, M. A., & Turner, C. R. (2016). The ecology of environmental DNA and im-plications for conservation genetics. Conservation Genetics, 17(1), 1–17. 10.1007/s10592-015-0775-4

55. Stewart, K. A. Understanding the Effects of Biotic and Abiotic Factors on Sources of Aquatic Environmental DNA. Biodivers. Conserv. 2019, 28(5), 983–1001. 10.1007/s10531-019-01709-8.

56. Li, J., Leavey, A., Wang, Y., O’Neil, C., Wallace, M. A., Burnham, C.-A. D., Boon, A. C., Bab-cock, H., & Biswas, P. (2018). Comparing the performance of 3 bioaerosol samplers for influenza virus. Journal of Aerosol Science, 115, 133–145. 10.1016/j.jaerosci.2017.08.007

57. Bessey, C.; Gao, Y.; Truong, Y. B.; Miller, H.; Jarman, S. N.; Berry, O. Comparison of Materials for Rapid Passive Collection of Environmental DNA. Mol. Ecol. Re-sour. 2022, 22(7), 2559–2572. 10.1111/1755-0998.13640.

58. Jane, S. F., Wilcox, T. M., McKelvey, K. S., Young, M. K., Schwartz, M. K., Lowe, W. H., Letcher, B. H., & Whiteley, A. R. (2015). Distance, flow and PCR inhibition: eDNA dynamics in two headwater streams. Molecular Ecology Resources, 15(1), 216–227. 10.1111/1755-0998.12285

59. Takahara, T.; Minamoto, T.; Yamanaka, H.; Doi, H.; Kawabata, Z. Estimation of Fish Biomass Using Environmental DNA. PLoS ONE 2012, 7(4), e35868. 10.1371/journal.pone.0035868.

60. Waza, A.; Schneiders, K.; May, J.; Rodríguez, S.; Epple, B.; Kandler, K. Field Comparison of Dry Deposition Samplers for Collection of Atmospheric Mineral Dust: Results from Single-Particle Characterization. Atmos. Meas. Tech. 2019, 12(12), 6647–6665. 10.5194/amt-12-6647-2019.

