## Supplementary Information for "A breath of fresh air: comparative evaluation of passive versus active airborne eDNA sampling strategies"

*Supplementary Methods – Nutshell eDNA sampler*

The Nutshell eDNA sampling device was developed with the goal to lower the barrier to collecting airborne eDNA. Its developer realized the relevance of airborne eDNA analysis to support a science-based understanding of biodiversity and ecosystem change but found himself dissatisfied with the technology available for this kind of work, which tends to be expensive, unwieldy, and in need of electrical power to operate. An affordable, easy to manufacture and operate, passive device should go a long way to increase the availability of airborne eDNA sampling for professional and non-professional biologists alike (e.g. for remote field work, or in a citizen science context). Additional design requirements were for the device to be easy to adapt to different sampling contexts and to be sustainable (i.e. made of recyclable materials and for repeated use) (Figure 1).

This resulted in a 10x10x20 cm device which consists of a Savonius-type wind turbine, a 19.6 mm diameter / 73 mm long open core to house any filter of choice, and a mounting part to fix the sampler to branch, post, or tripod, using zip-ties. The theory of operation for this device is that energetic, particle-laden air drives the turbine, which makes the air lose energy in the process. The air slows further down in the open core, which initiates a sedimentation process that settles DNA-laden particles in the filter. The core has an exit at the underside for easier passage of air through the device. Additionally, the turbine acts as a housing to protect the DNA in the filter from UV light, moisture, and interference from macroscopic animals.

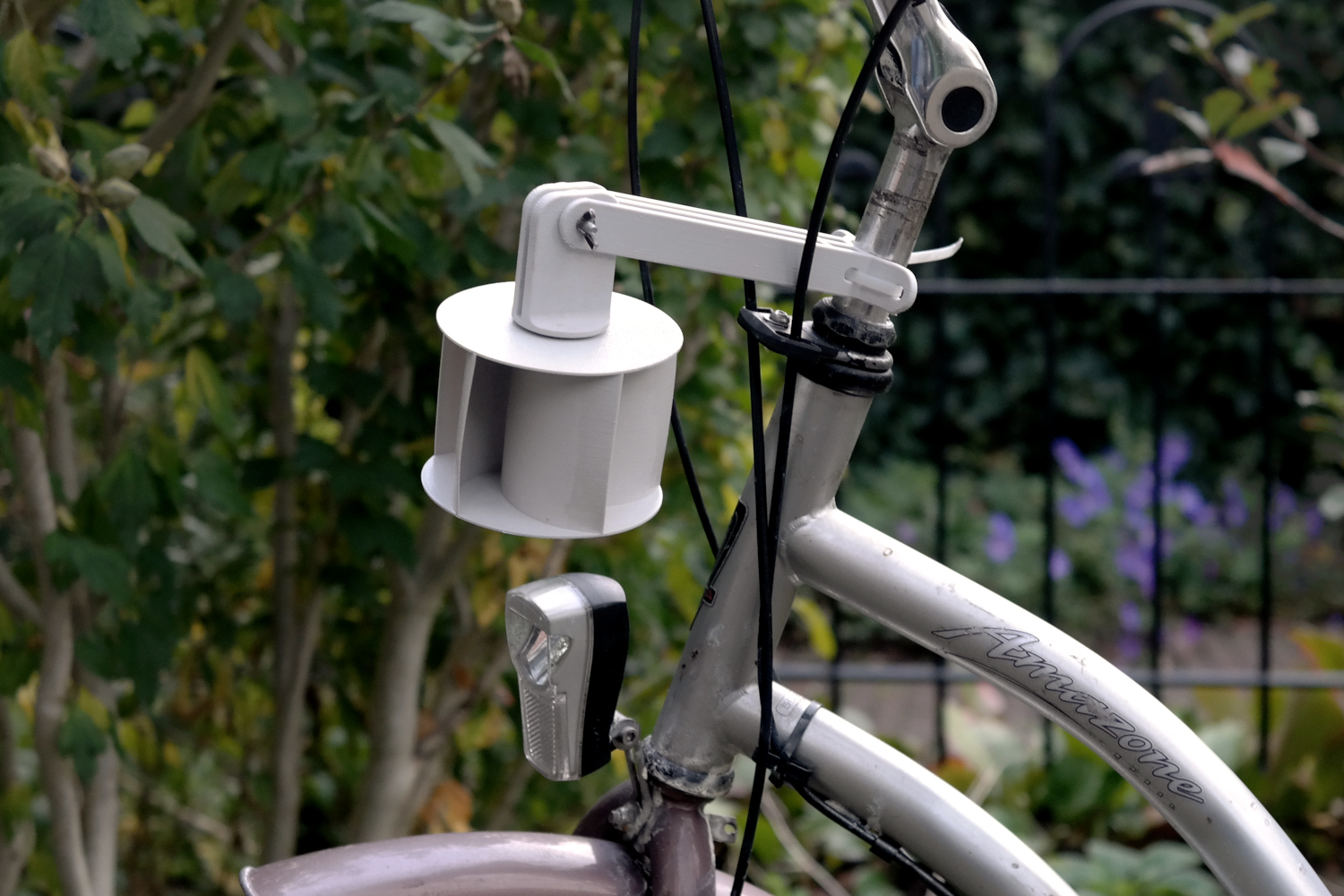

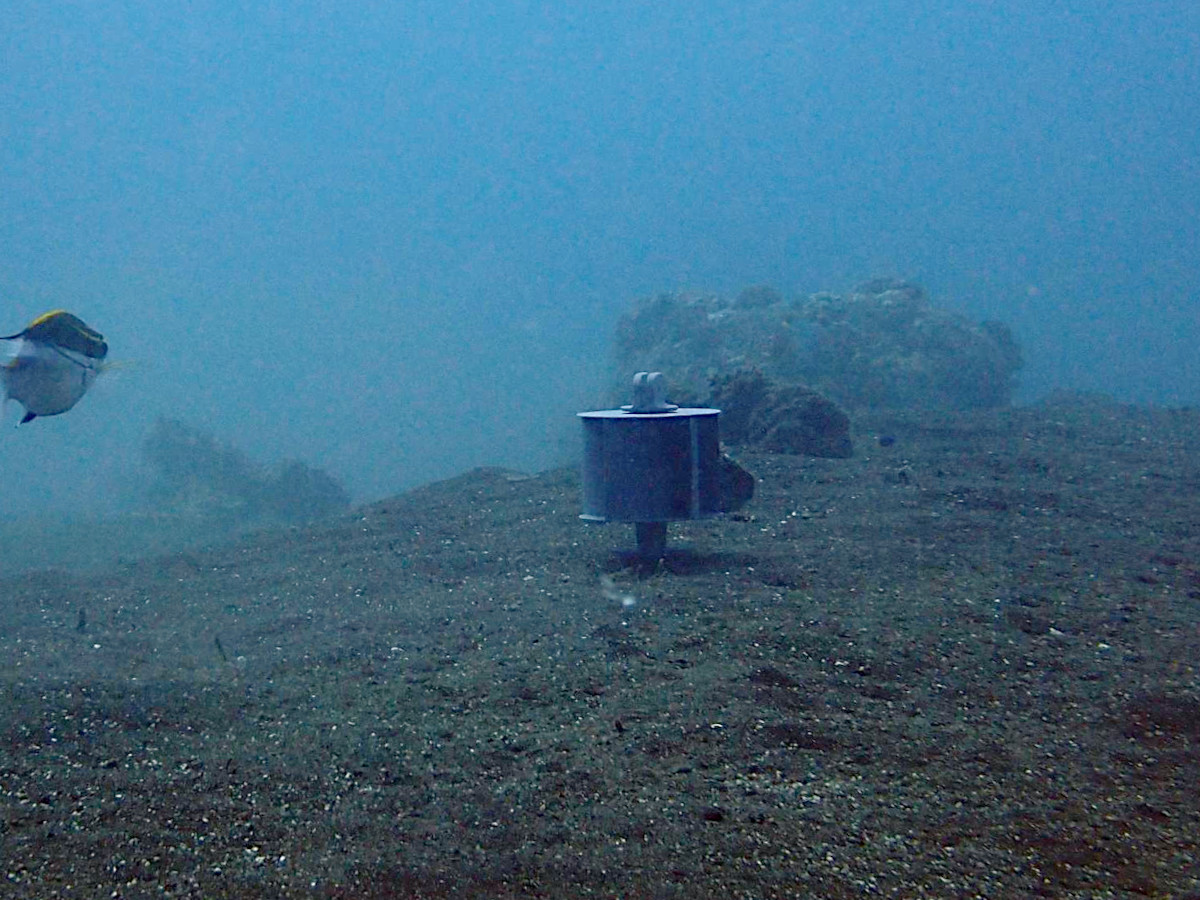

**Figure S1:** Nutshell eDNA sampler adapted for mobile (left) and marine (right) eDNA collection.

Most of the parts can be 3D printed on any off-the-shelf fused-filament 3D printer, the developer currently uses a Bambulab P1P. Most parts are printed at a layer height of 0.2 mm, except for the threaded parts which need to be printed at a higher 0.12 mm resolution for the threads to function. All filament-based parts are printed with a fill of 30%, using PETG filament. PETG is relatively hard and tough, has great layer adhesion, can be recycled, and is UV-resistant enough for outdoors use. The geometry of the core part makes it hard to print on a filament printer, so for this part a stereolithography printer is used instead, an Elegoo Saturn 4 Ultra with Anycubic tough resin 2.0.

Material per sampler comes down to 26.30 m of 1.75 mm diameter PETG filament, 4.15 ml tough resin, two POM6804 plastic and glass ball bearings, a m4/30 mm wingbolt, a m4 wingnut, cyanoacrylate (´super´) glue to fix the ball bearings to the sampler, MEK solvent to chemically weld some of the printed parts together, and a zip-tie to mount the device to a support structure (Figure 2). At the time of writing, total material cost is around 4 Euro per sampler. All non-printed parts are standard and easily available. Instructions for construction and other additional information can be found at [1].

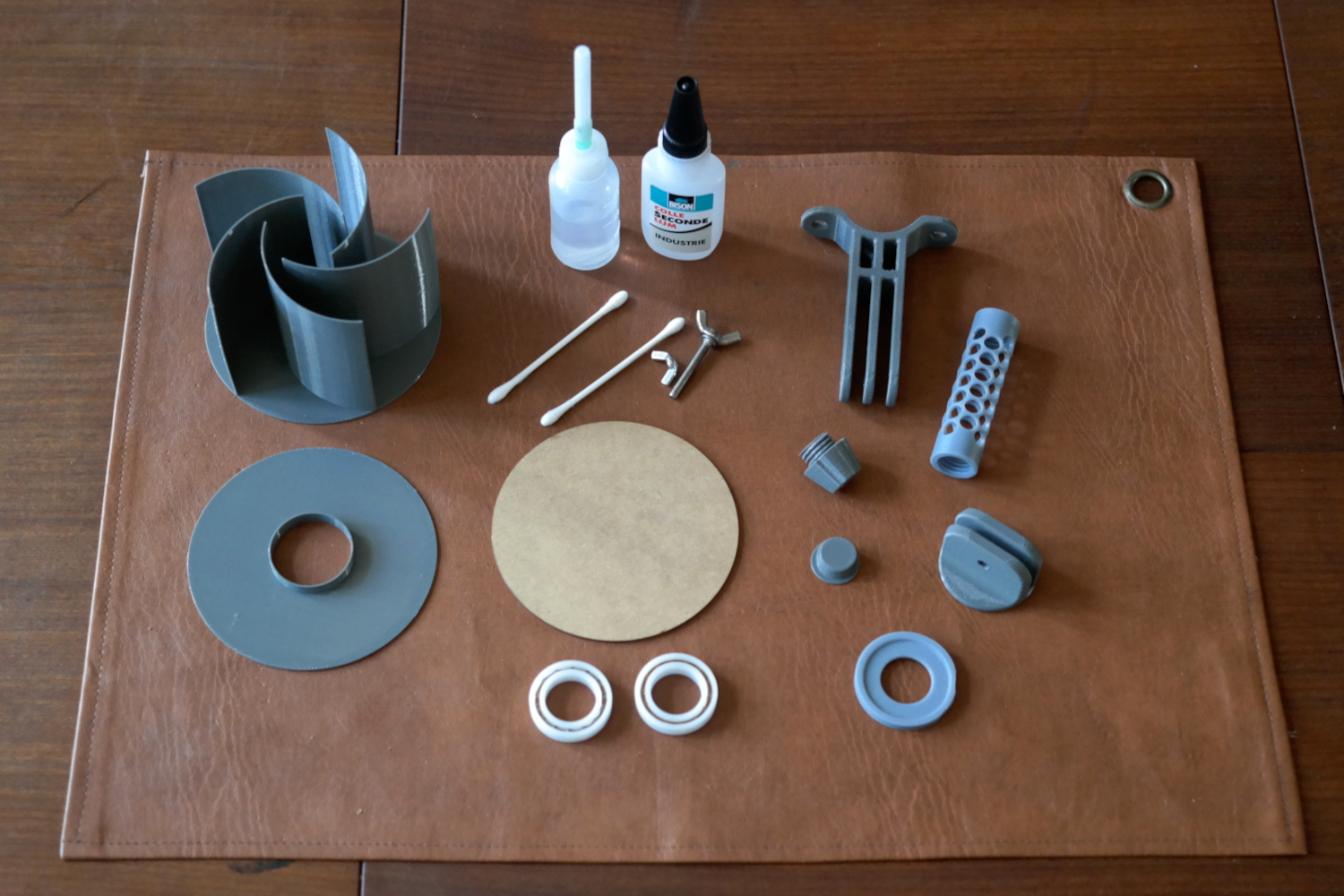

**Figure S2:** Parts and tools needed to assemble the Nutshell eDNA sampler.

The Nutshell eDNA sampler is open source and published on a GitHub page since August 24, 2023, under a CERN Open Hardware License, Version 2 - Weakly Reciprocal [2]. In broad strokes, this means that anyone may print, use, adapt, and further distribute this device, provided that the accompanying information and license is distributed along with it. Additionally, any adaptations to the design have to be made publicly available, under the same license. Open sourcing the design was done to facilitate the spread and use of this device, with an eye on future collaborative development and building a user community. The developer is happy to collaborate on projects involving this sampler, contact information is provided on the referenced GitHub page [1].

**References**

[1] Luursema, J. M. (2023). Nutshell eDNA-sampler. Available at [https://github.com/J4n-M44rt3n/](https://github.com/J4n-M44rt3n/DNAir-sampler)Nutshell-eDNA[-sampler](https://github.com/J4n-M44rt3n/DNAir-sampler) [accessed September 25, 2025].

[2] CERN (2020). CERN Open Hardware License V2. Available at <https://gitlab.com/ohwr/project/cernohl/-/wikis/uploads/f773df342791cc55b35ac4f907c78602/cern_ohl_w_v2.pdf>

TABLE S1 Sampling schedule across the three weeks of sampling in Rotterdam Zoo. P-’number’ indicates the collection of a Nutshell eDNA sample for each of the six locations, with the number indicating the number of hours the sampler had been hung at its location before collection. P-neg is a field-negative Nutshell eDNA-sample.

|  | Monday | Tuesday | Wednesday | Thursday | Friday | Saturday |
| --- | --- | --- | --- | --- | --- | --- |
| Week 1 | **25^th^ of March 2024**  10:30  Hung passive samplers on assigned locations  13:30  Active air eDNA sampling on assigned locations  16:30  P-6 collection | **26^th^ of March 2024**  10:30  P-24 collection | **27^th^ of March 2024**  10:30  P-48 collection | **28^th^ of March 2024**  10:30  P-72 collection | **29^th^ of March 2024**  10:30  P-96 collection  13:00  P-neg collection | **30^th^ of March 2024** |
| Week 2 |  | **2^nd^ of April 2024**  9:00  Hung passive samplers on assigned locations  11:30  Active air eDNA sampling on assigned locations  15:00  P-6 collection | **3^rd^ of April 2024**  9:00  P-24 collection | **4^th^ of April 2024**  9:00  P-48 collection | **5^th^ of April 2024**  9:00  P-72 collection | **6^th^ of April 2024**  9:00  P-96 collection  11:30  P-neg collection |
| Week 3 | **8^th^ of April**  9:00  Hung passive samplers on assigned locations  11:30  Active air eDNA sampling on assigned locations  15:00  P-6 collection | **9^th^ of April**  9:00  P-24 collection | **10^th^ of April**  9:00  P-48 collection | **11^th^ of April**  9:00  P-72 collection | **12^th^ of April**  9:00  P-96 collection  11:30  P-neg collection | **13^th^ of April** |

TABLE S2 Total number of individuals (Ind.) housed within the Rotterdam Zoo at the time of sampling for each detected zoo species, the average mass for the detected zoo species, with affiliated reference. Asterix denotes outliers based on total read counts >35000 sequences.

| Species | Ind. | Average mass per individual (kg) | Total Biomass (kg) | Source |
| --- | --- | --- | --- | --- |
| *Antilope cervicapra* | 17 | 36.5 | 620.5 | <https://animaldiversity.org/accounts/Antilope_cervicapra/> |
| *Bison bison* | 12 | 609 | 7308 | <https://animaldiversity.org/accounts/Bison_bison/> |
| *Callosciurus prevostii* | 2 | 0.45 | 0.9 | <https://animaldiversity.org/accounts/Callosciurus_prevostii/> |
| **Camelus bactrianus* | 4 | 708 | 2832 | <https://a-z-animals.com/animals/bactrian-camel/> |
| *Chauna torquata* | 2 | 3 | 6 | <https://www.zooparc.nl/dier/kuifhoenderkoet> |
| *Diceros bicornis* | 3 | 1100 | 3300 | <https://animaldiversity.org/accounts/Diceros_bicornis/> |
| **Elephas maximus* | 4 | 3750 | 15000 | <https://animaldiversity.org/accounts/Elephas_maximus/> |
| *Equus quagga* | 3 | 312.5 | 937.5 | <https://a-z-animals.com/animals/zebra/> |
| **Giraffa camelopardalis* | 8 | 1240 | 9920 | [https://a-z-animals.com/animals/giraffe/](https://animaldiversity.org/accounts/Haliaeetus_pelagicus/) |
| *Haliaeetus pelagicus* | 2 | 7.5 | 15 | https://animaldiversity.org/accounts/Haliaeetus_pelagicus/ |
| *Lutra lutra* | 3 | 6.75 | 20.25 | https://animaldiversity.org/accounts/Lutra_lutra/ |
| *Macaca silenus* | 5 | 6.5 | 32.5 | <https://animaldiversity.org/accounts/Macaca_silenus/> |
| *Mareca strepera* | 48 | 0.765 | 36.72 | <https://www.wildlifetrusts.org/wildlife-explorer/birds/waterfowl/gadwall> |
| *Merops nubicus* | 8 | 0,0465 | 0.372 | <https://animaldiversity.org/accounts/Merops_nubicus/> |
| *Nanger dama* | 8 | 55 | 440 | <https://animaldiversity.org/accounts/Nanger_dama/> |
| *Nycticorax nycticorax* | 7 | 0.8 | 5.6 | <https://animaldiversity.org/accounts/Nycticorax_nycticorax/> |
| *Pelecanus crispus* | 38 | 11.5 | 437 | <https://galicica.org.mk/en/gallery/the-dalmatian-pelican-pelecanus-crispus/> |
| *Phoenicopterus roseus* | 146 | 2.45 | 357.7 | <https://animaldiversity.org/accounts/Phoenicopterus_roseus/> |
| *Rhinoceros unicornis* | 1 | 1950 | 1950 | <https://animaldiversity.org/accounts/Rhinoceros_unicornis/> |
| *Struthio camelus* | 2 | 110 | 220 | <https://animaldiversity.org/accounts/Struthio_camelus/> |
| *Tadorna tadorna* | 16 | 1.1 | 17.6 | <https://www.wildlifetrusts.org/wildlife-explorer/birds/waterfowl/shelduck> |
| *Theropithecus gelada* | 17 | 17 | 289 | <https://animaldiversity.org/accounts/Theropithecus_gelada/> |
| *Trachypithecus francoisi* | 4 | 9.15 | 36.6 | <https://animaldiversity.org/accounts/Trachypithecus_francoisi/> |
| **Vulpes lagopus* | 5 | 5.2 | 26 | <https://animaldiversity.org/accounts/Vulpes_lagopus/> |

TABLE S3 Table including the total read counts across all samples for each identified species per primer set and combined. Species highlighted in grey are zoo residents.

|  |  | Hits per primer set | |  |
| --- | --- | --- | --- | --- |
| Species | Common name | 12S | 16S | Total read count |
| *Accipiter nisus* | Eurasian sparrowhawk | 4690 | 0 | 4690 |
| *Alopochen aegyptiaca* | Egyptian goose | 7108 | 0 | 7108 |
| *Alytes obstetricans* | Common midwife toad | 14369 | 706 | 15075 |
| *Alytes muletensis* | Majorcan midwife toad | 75664 | 2974 | 78638 |
| *Anas platyrhynchos* | Mallard | 21527 | 0 | 21527 |
| *Antilope cervicapra* | Indian antelope | 0 | 197 | 197 |
| *Apodemus sylvaticus* | Wood mouse | 33 | 1250 | 1283 |
| *Ardea cinerea* | Grey heron | 2457 | 0 | 2457 |
| *Arvicola amphibius* | European water vole | 0 | 312 | 312 |
| *Aythya sp.* | Pochard species | 179 | 0 | 179 |
| *Bison bison* | American bison | 0 | 469 | 469 |
| *Bombina orientalis* | Oriental fire-bellied toad | 4225 | 516 | 4741 |
| *Bos taurus* | Domestic cow | 8367 | 141589 | 149956 |
| *Branta nigricans orientalis* | Brent goose | 10577 | 0 | 10577 |
| *Bufo bufo* | Common toad | 323 | 145 | 468 |
| *Callosciurus prevostii* | Prevost's squirrel | 0 | 301 | 301 |
| *Camelus bactrianus* | Bactrian camel | 13090 | 31539 | 44629 |
| *Canis lupus familiaris* | Dog | 13 | 31175 | 31188 |
| *Capra hircus* | Domestic goat | 0 | 338 | 338 |
| *Capra pyrenaica* | Iberian ibex | 0 | 726 | 726 |
| *Cavia porcellus* | Guinea pig | 203 | 228 | 431 |
| *Cervus elaphus* | Red deer | 1768 | 43 | 1811 |
| *Chauna torquata* | Southern screamer | 717 | 0 | 717 |
| *Clupea harengus* | Atlantic herring | 0 | 7 | 7 |
| *Columba livia* | Rock dove | 65374 | 0 | 65374 |
| *Columba oenas* | Stock dove | 1141 | 0 | 1141 |
| *Corvus splendens* | House crow | 4338 | 0 | 4338 |
| *Crocidura russula* | Greater white-toothed shrew | 0 | 228 | 228 |
| *Cyprinus carpio* | Common carp | 110993 | 2138 | 113131 |
| *Diceros bicornis* | Black rhinoceros | 4147 | 2086 | 6233 |
| *Elephas maximus* | Asian elephant | 0 | 36984 | 36984 |
| *Equus caballus* | Horse | 0 | 2082 | 2082 |
| *Equus quagga* | Plains zebra | 0 | 1076 | 1076 |
| *Felis catus* | House cat | 1405 | 13776 | 15181 |
| *Fringilla coelebs* | Eurasian chaffinch | 5306 | 0 | 5306 |
| *Fulica atra* | Eurasian coot | 62 | 0 | 62 |
| *Gallinula chloropus* | Common moorhen | 3680 | 0 | 3680 |
| *Gallus gallus* | Domestic fowl | 30780 | 0 | 30780 |
| *Giraffa camelopardalis* | Giraffe | 20085 | 26044 | 46129 |
| *Gobio gobio* | Gudgeon | 1131 | 44 | 1175 |
| *Haliaeetus pelagicus* | Steller's sea eagle | 1195 | 0 | 1195 |
| *Ichthyosaura alpestris* | Alpine newt | 382 | 189 | 571 |
| *Lama guanicoe* | Guanaco | 0 | 132 | 132 |
| *Larus marinus* | great black-backed gull | 496 | 0 | 496 |
| *Lepus europaeus* | European hare | 0 | 1289 | 1289 |
| *Leuciscus aspius* | Asp | 4824 | 0 | 4824 |
| *Lissotriton helveticus* | Palmate newt | 473 | 0 | 473 |
| *Lutra lutra* | European otter | 20 | 1111 | 1131 |
| *Macaca silenus* | Lion-tailed macaque | 0 | 79 | 79 |
| *Mallotus villosus* | Capelin | 2605 | 588 | 3193 |
| *Mareca strepera* | Gadwall | 3014 | 0 | 3014 |
| *Merops nubicus* | Northern carmine bee-eater | 20 | 0 | 20 |
| *Microtus arvalis* | Common vole | 0 | 582 | 582 |
| *Mus musculus* | House mouse | 676 | 725 | 1401 |
| *Nanger dama* | Addra gazelle | 0 | 329 | 329 |
| *Nycticorax nycticorax* | Black-crowned night heron | 1816 | 0 | 1816 |
| *Oncorhynchus clarkii lewisi* | Westslope cutthroat trout | 736 | 0 | 736 |
| *Oncorhynchus kawamurae* | Black kokanee | 2383 | 0 | 2383 |
| *Oryctolagus cuniculus* | European rabbit | 0 | 2725 | 2725 |
| *Osmerus eperlanus* | European smelt | 677 | 13 | 690 |
| *Ovis aries* | Domestic sheep | 0 | 2079 | 2079 |
| *Passer domesticus* | House sparrow | 485 | 0 | 485 |
| *Pelecanus crispus* | Dalmatian pelican | 171 | 0 | 171 |
| *Pelophylax lessonae* | Pool frog | 7780 | 0 | 7780 |
| *Phasianus colchicus* | Ring-necked pheasant | 229 | 0 | 229 |
| *Phoenicopterus roseus* | Greater flamingo | 629 | 0 | 629 |
| *Phoxinus phoxinus* | Common minnow | 2579 | 0 | 2579 |
| *Pterodroma heraldica* | Herald petrel | 3 | 0 | 3 |
| *Rana temporaria* | Common frog | 2210 | 0 | 2210 |
| *Rattus norvegicus* | Brown rat | 0 | 3329 | 3329 |
| *Rhinoceros unicornis* | Indian rhinoceros | 255 | 6340 | 6595 |
| *Rutilus rutilus* | Common roach | 0 | 182 | 182 |
| *Salmo Salar* | Atlantic salmon | 1665 | 0 | 1665 |
| *Scomber colias* | Atlantic chub mackerel | 667 | 0 | 667 |
| *Solea solea* | Common sole | 1017 | 0 | 1017 |
| *Sprattus sprattus* | European sprat | 12002 | 0 | 12002 |
| *Squalius cephalus* | Common chub | 763 | 28 | 791 |
| *Struthio camelus* | Common ostrich | 1381 | 0 | 1381 |
| *Sturnus vulgaris* | Common starling | 6728 | 0 | 6728 |
| *Sus scrofa* | Domestic pig | 2985 | 2023 | 5008 |
| *Tadorna tadorna* | Common shelduck | 1056 | 0 | 1056 |
| *Theropithecus gelada* | Gelada | 0 | 721 | 721 |
| *Threskiornis aethiopicus* | African sacred ibis | 8 | 0 | 8 |
| *Trachypithecus francoisi* | Francois' leaf monkey | 0 | 3 | 3 |
| *Triturus carnifex* | Italian crested newt | 3381 | 0 | 3381 |
| *Triturus karelinii* | Southern crested newt | 0 | 243 | 243 |
| *Turdus merula* | Common blackbird | 3796 | 0 | 3796 |
| *Vulpes lagopus* | Arctic fox | 1139 | 38566 | 39705 |

TABLE S4 Table showing the average distance between detected zoo species and the outdoor sampling location at which they were detected for each air eDNA sampler, including the average detection distance for zoo species across the different outdoor sampling locations per air eDNA sampler.

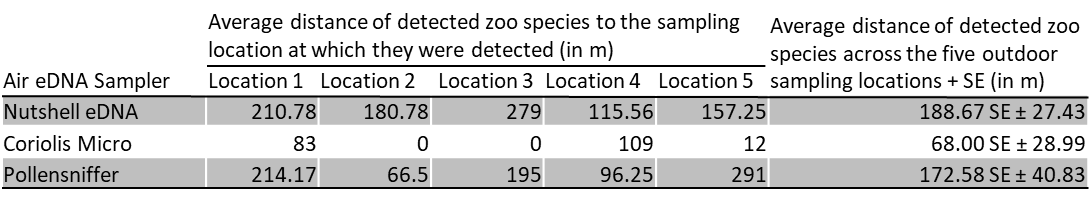

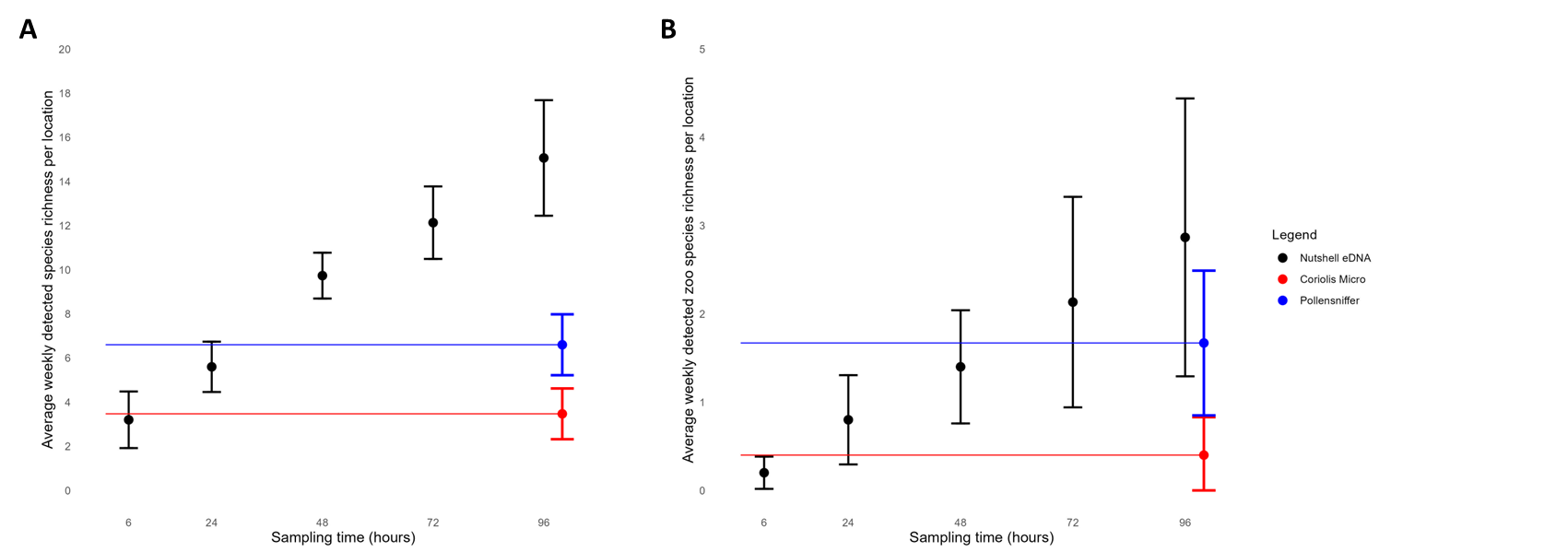

**Figure S3**: Species richness means (±SD) for outdoor sampling locations (sites 1-5). **(A)** Average weekly detected species richness per outdoor sampling location by the Nutshell eDNA sampler strategy (black) over 96 hours, juxtaposed against the mean (±SD) average weekly detected species richness per outdoor sampling location by the Coriolis Micro (red) and Pollensniffer (blue) static samples, with **(B)** showing total zoo resident species richness detection.

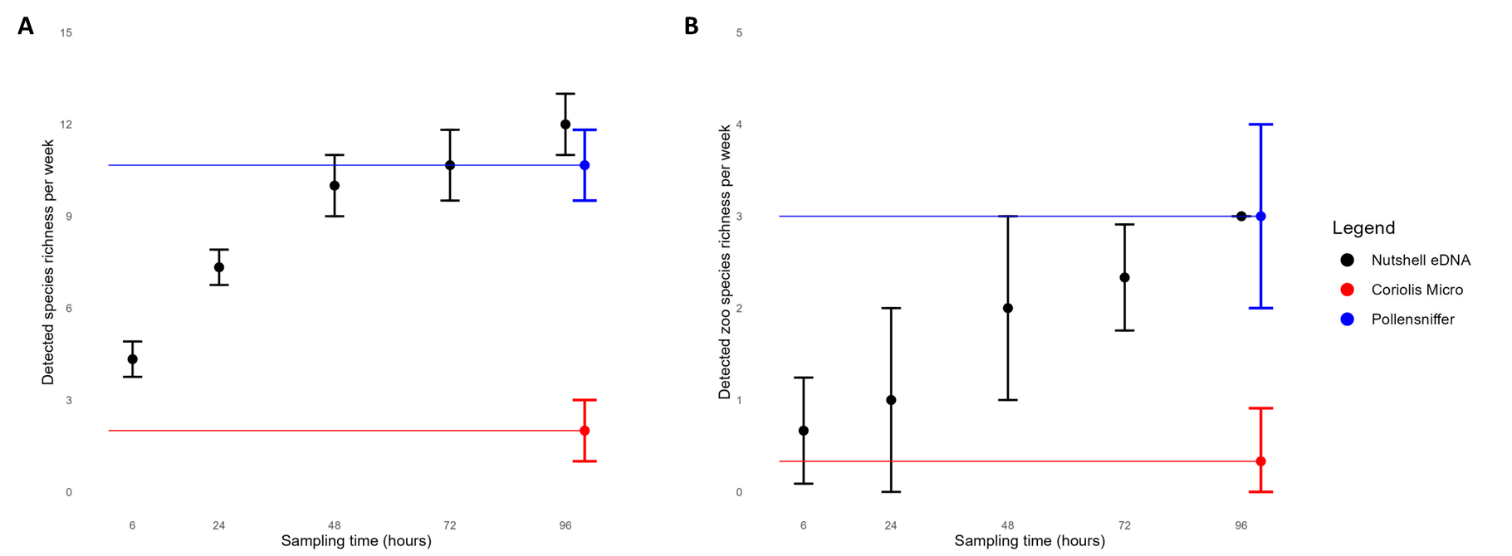

.

FIGURE S4 Species richness means (±SD) for the indoor sampling site (site 6) averaged across the three-week sampling campaign. **(A)** Average weekly detected species richness by the Nutshell eDNA sampler strategy (black) over 96 hours, juxtaposed against the mean (±SD) average detected species richness by the Coriolis Micro (red) and Pollensniffer (blue) static samples per week (in triplicate), with **(B)** showing total zoo resident species richness detection.

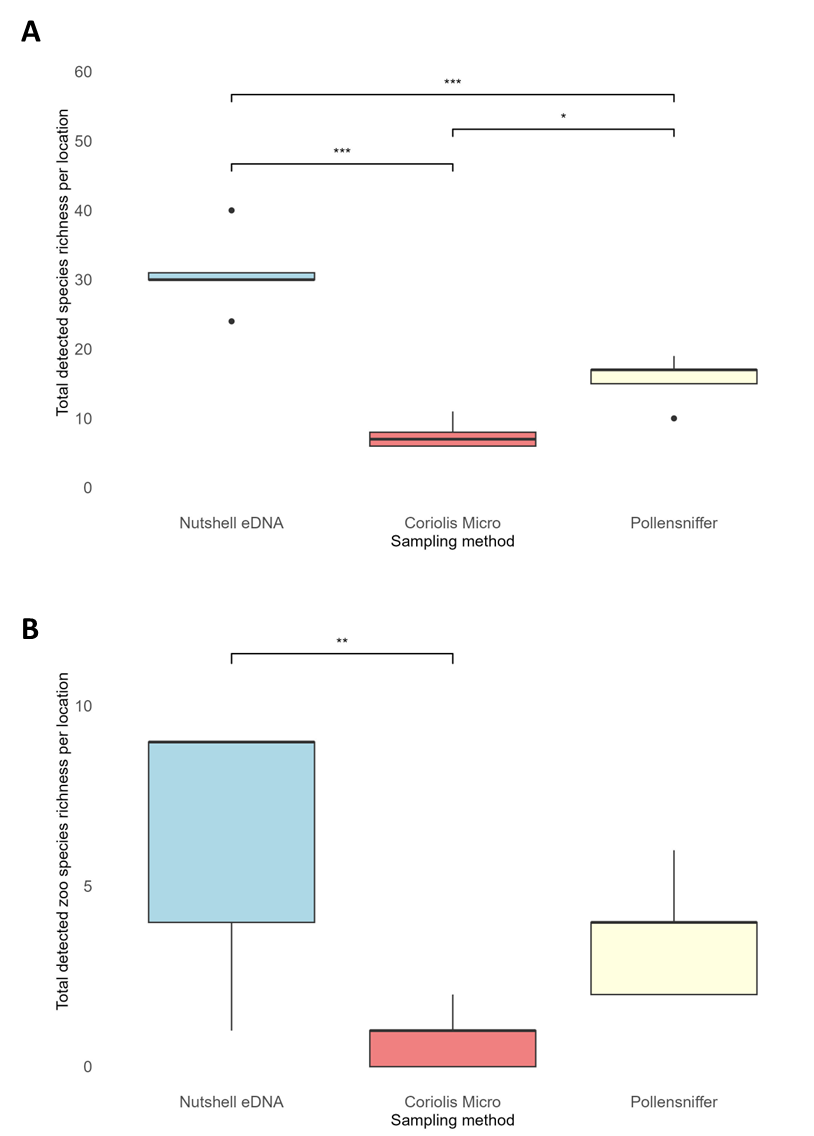

FIGURE S5 Boxplots showing the total detected species richness (A) and total detected zoo species richness (B) per outdoor sampling location for each of the air eDNA samplers.

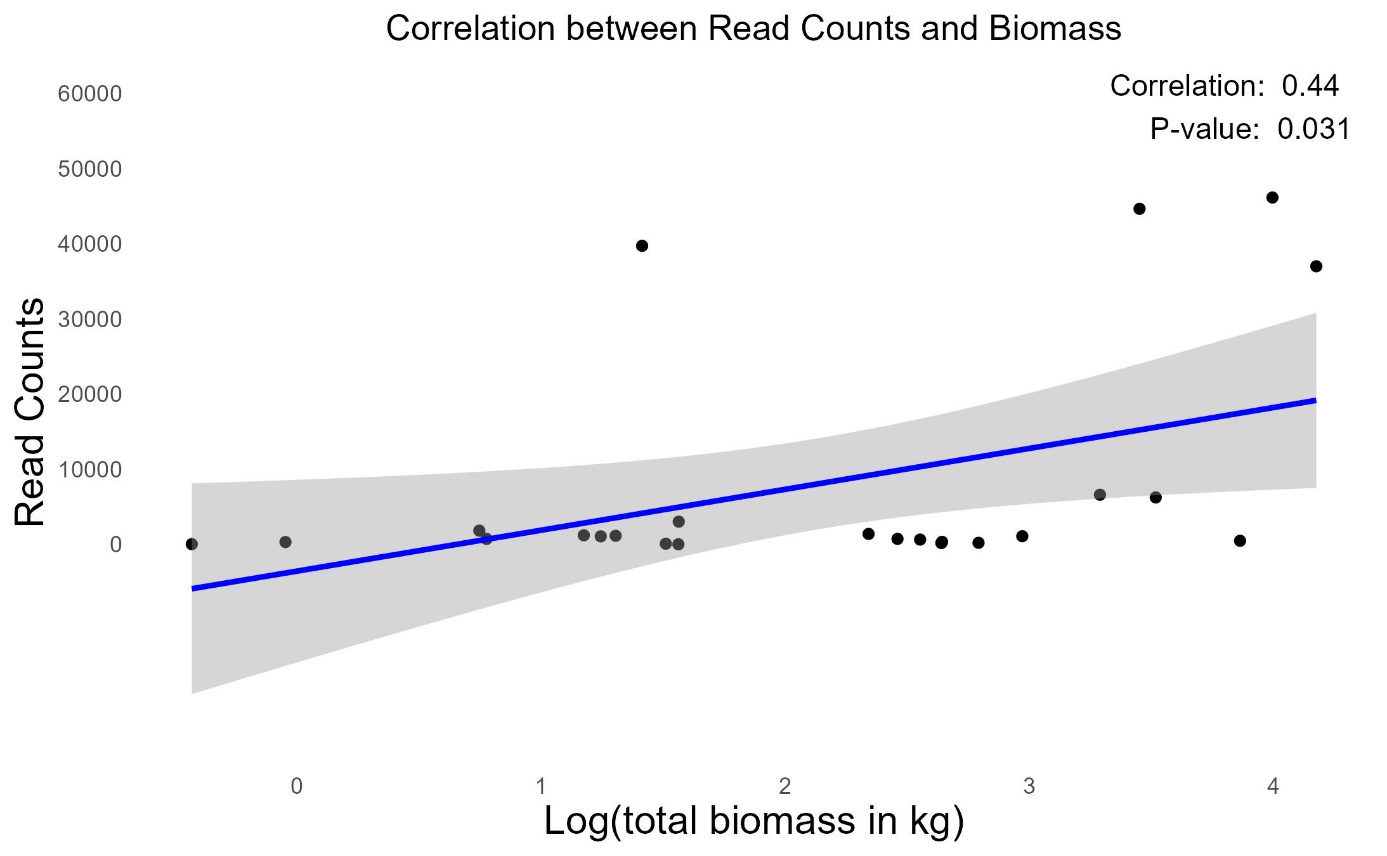

**a**

**b**

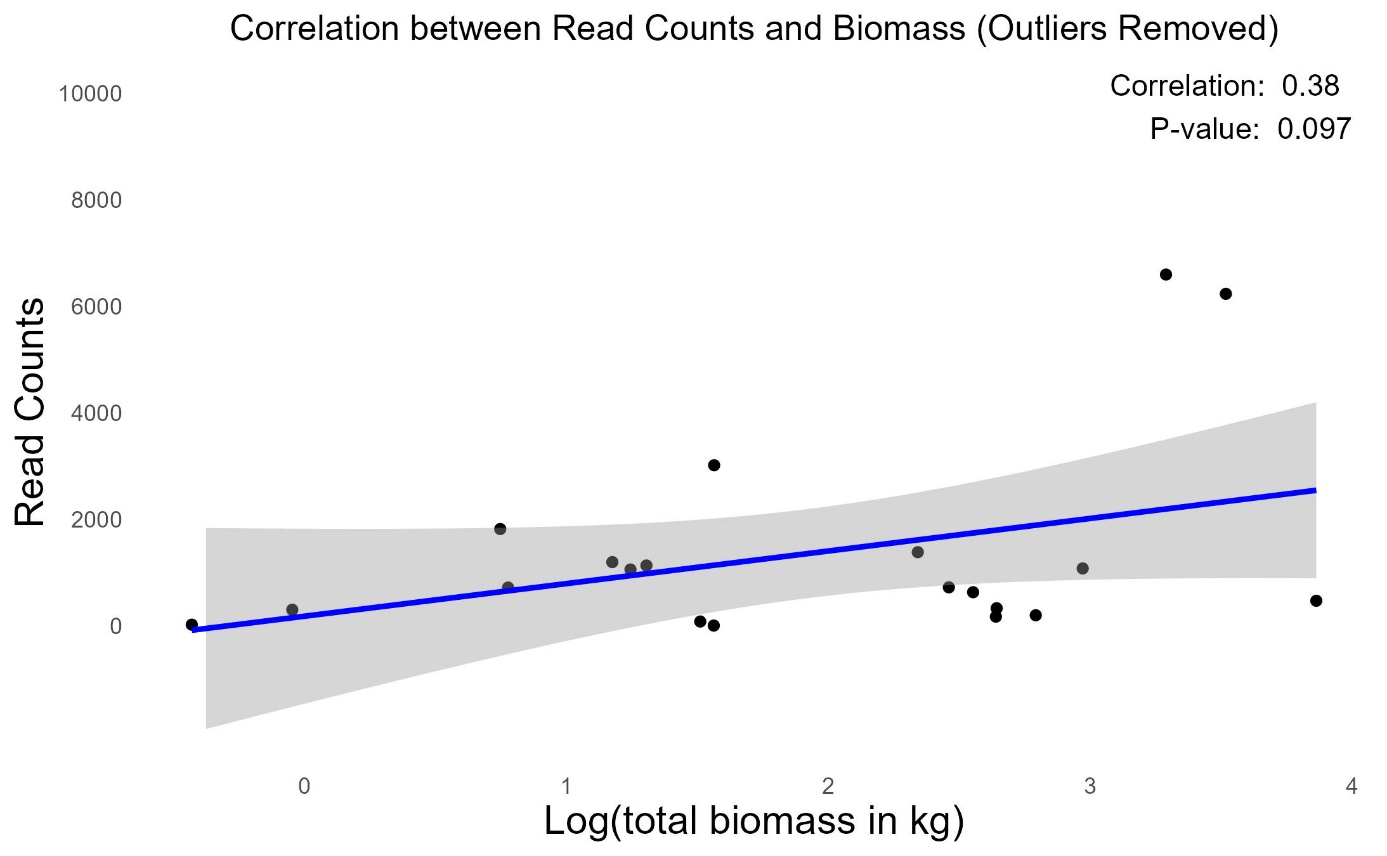

FIGURE S6 Correlation graph between total sequencing reads (read counts) and Log transformed total biomass (kg) of detected zoo species (A), and with 4 outliers removed (B).
